# Disentangling different sources of variation in functional responses: between-individual variability, measurement error and inherent stochasticity of the prey-predator interaction process

**DOI:** 10.1101/2025.07.09.663660

**Authors:** Charlotte Baey, Sylvain Billiard, Maud Delattre

## Abstract

The consumption rate of prey by predators, or functional responses, are known to be highly variable even within a single population. Identifying and estimating the different sources of variation of functional responses is a long-standing challenge. We develop here a statistical framework derived from a mechanistic stochastic process model that explicitly accounts for different sources of variation. We apply it to disentangle and estimate in particular 1) residual variance due to measurement errors and model misspecification, 2) between-predator variability, and 3) the *interaction stochasticity, i.e*. the intrinsic and mechanistic variability due to interactions processes between prey and predators. We show that it is possible to estimate these sources of variation under realistic experimental conditions. Our results also show that model fitting can compensate by overestimating residual source of variation, leading to biased parameter estimates when interaction stochasticity is misspecified. Applied to empirical data, the model reveals that standard assumptions, such as prey renewal and lack of spatial structure, fail to capture observed variability. We also show how experimental design affects parameter identifiability, highlighting the trade-off between the number of individuals and repeated observations.

## 1 Introduction

Functional responses are functions describing the rate of interactions between two or more species in population or community ecology models. For example, functional responses relate the consumption rate of prey by a predator and the prey density in the environment, such as captured by the classical Holling types I, II or III models or many others [16, 17]. As prey and predator dynamics are coupled through such a functional response, the chosen functional form introduced into models generally has a dramatic impact on the conditions for the coexistence between species, the stability of a community and the speed of the flux of matter and energy in an ecosystem. As a consequence, it is a major challenge in ecology to properly estimate functional responses and their parameters from data, as shown by the large number of experiments devoted to this goal that have been conducted in the last decades [17, 27, 41].

It has long been recognized that prey consumption rates measured in experiments typically show unexpected large variance as well as heteroscedasticity [38]. For instance, under the assumption that the number of prey eaten by a predator follows a binomial distribution, data typically show overdispersion [38, 23, 7, 14]. A beta-binomial distribution was then proposed as an alternative for inference as it allows for overdispersion, albeit at the cost of a higher number of parameters to be estimated [7, 14]. However, no biological justification was given for such a choice. More recently, consumption rates have been shown to commonly have a standard deviation of the same order of magnitude than the mean [4, 31]. Such a large variance in data raises the challenge to correctly identify and quantify the possible sources of such a large variability.

Among all possible sources of variation of functional responses, as observed in experiments or natural populations, four in particular are the focus of identification and estimation: i) residual variance due to measurements errors (*e*.*g*. in the initial number of prey provided to a predator, or in the number of prey recorded as eaten when they were in fact removed from the experiment for other reasons [18, 3]) or model misspecification (*e*.*g*. when some processes are neglected such as between-predator competition); ii) between-individual variability in predators’ or prey’ traits involved in their interactions (*e*.*g*. individuals size, personality, age or satiety [23, 26, 30, 34]); iii) variability of the environment (*e*.*g*. dimension, temperature, vegetative cover [26, 39]); iv) *interaction stochasticity, i*.*e*. a mechanistic intrinsic source of variation due to the foraging and the interaction processes themselves [6, 4, 12, 31], (*e*.*g*. because finding a prey, capturing it and succeeding in feeding on it are random events).

These different sources of variability of functional responses are, at most, partially taken into account, when they are. Functional responses are generally estimated by fitting non-linear models that describe the relationship between prey density and consumption rate to data collected in a controlled environment where the number of prey consumed per unit of time per predator are measured at different prey densities (*e*.*g*. [3, 32], *see also [27, 41] for extended reviews), rarely in natural populations (e*.*g*. [26, 33]). Several methodological approaches are applied to select which functional response best fits the data among possible alternatives, or to estimate their parameters. Depending on the chosen approach, different sources of variations of the functional response could, or could not, be evaluated.

A first approach consists in estimating parameters using independent behavioral measurements and direct observations, for example for the time taken to handle a prey, or the movement speed, and plug these estimates in candidate functions, in order to predict consumption rates, and sometimes to qualitatively compare predictions to data (e.g. [23, 37, 3, 22, 5]).

A second approach is based on directly fitting deterministic equations to observed prey consumption rates, or the number of prey consumed after a given fixed time, using least squares or likelihood optimization procedures [15, 38, 28, 8, 39, 35, 37]. In the first and second approaches, variability of the functional responses and its possible sources are totally neglected.

Third, goodness of fit or regression procedures are applied to estimate the parameter of a deterministic equation specifying a particular error structure with prey density as explanatory variable [38, 7, 37, 33, 14, 32], sometimes associated with other independent variables such as environmental treatments [24]. In this case, the only quantified source of variability are the measurements error and model misspecification estimated as the residual variance.

Fourth, by using a maximum-likelihood procedure, likelihood ratio test, [6] fitted a Holling II-type model including both residual variance and interaction stochasticity, but neglecting between-predator traits variability, with data lacking repeated measures on identical predator individuals, precluding a correct estimate of the various sources of variation. Finally, a last approach consists in directly including inter-individual variability in predators into non-linear regression models by adding individuals as a random effect, or considering traits covariates such as social status [26]. In such an approach, measurements errors and variations among predators are thus evaluated as parameters of the supposed error structure (residual variance) or the distribution of the random effect (referred to hereafter as between-individual predators variance).

Between-predator variability is sometimes indirectly evaluated in successive steps: the parameters of the functional response are first estimated from direct observations data (as in the first approach, see before), or from within-individual fitting of functional responses (as in the second or third approaches, see before). Correlations are then searched between these estimated individual parameters with covariates *(e*.*g*. individuals size or behavior), the environment (*e*.*g*. temperature), either for different individuals of a single species [34], or for different individuals for different species [40, 30], or through comparison between the variance observed in data and the one predicted in a partial differential equation model accounting for inter-individual variance only [23].

In summary, to our knowledge, no existing statistical framework for inferring functional responses adequately accounts for the various sources of variation described above, namely those generally considered as the most relevant. It is yet important as model selection and parameter estimations of functional responses are necessarily biased or lack precision because of such a large uncontrolled and unidentified variance [27], for three main reasons.

First because it is common that several different candidate non-linear models fit close enough to the data so that it is impossible to select one among others, because of the nature of data and non-linear models themselves (*e*.*g*. [3, 35, 27]). Second, if the between-individual variability in predators or the interaction stochasticity are neglected, they are necessarily conflated with residual noise, leading to poor parameter estimates. Third, because heteroscedasticity —unequal variation in errors across prey densities— is frequently observed. At low prey densities, errors may be relatively small, while at higher densities, variability in consumption rates often increases. Failing to account for these differences in error can result in biased estimates, particularly regarding the effect of prey density and other mechanistic parameters. For instance, if the main objective of an experiment is to test whether variability among predators is a major factor affecting functional responses, this variability must be properly isolated from other possible sources of variation. Otherwise, there is a risk of drawing incorrect conclusions. Therefore, it is essential to adopt a rigorous approach that accounts for these factors, not only to improve the accuracy of parameter estimates but also to ensure that the results can be interpreted and generalized to broader ecological contexts.

This paper aims to demonstrate how to conduct a robust statistical analysis of experimental data on functional responses, incorporating all the above mentionned sources of variability, in a one-step procedure (Fig. 1). It will illustrate how appropriate statistical modeling, combined with robust inference methods, can correct these biases. In particular, the use of mixed-effects models allows for inter-individual variability in predators to be accounted for, thereby improving the quality of parameter estimates. Our statistical framework will also be based on the elaboration of functional responses derived from stochastic processes modeling of the interaction between prey and predators, which is necessary to take into account variations due to interaction stochasticity from foraging and feeding processes themselves. We will describe the algorithmic procedure that allows estimation of such functional responses based on Stochastic Gradient Descent (SGD). We will then show how to account for the different possible sources of variation. We will also compare the reliability of the estimations of different experimental designs in order to propose guidelines for experimenters willing to estimate different sources of variation. Finally, as an important objective in the ecological literature is to find the functional response that best fits to data, we will illustrate how our statistical model can be used for model selection. As a proof of concept, we will compare models with or without the interaction stochasticity as a source of variability. Such a model comparison will allow to evaluate the relative importance of interaction stochasticity comparatively to between-predator variability and to residual variance.

**Figure 1:**
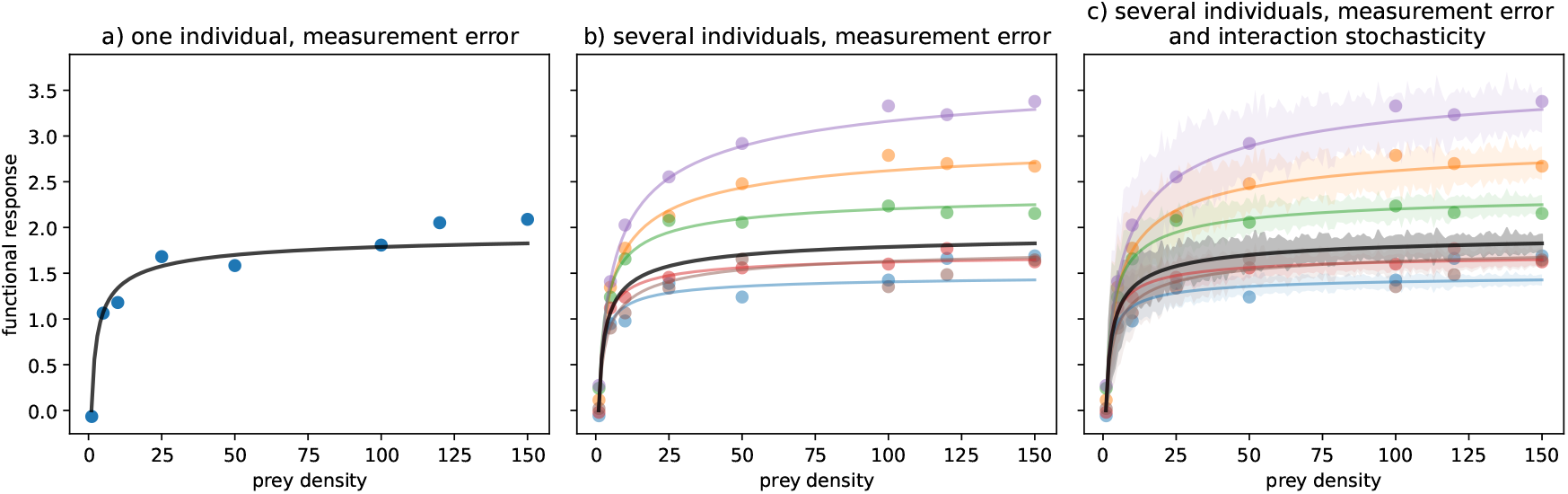
The different sources of variability in a functional response. In all plots, the black curve represents the mean functional response *r*. On the left, the dots corresponds to the mean functional response plus a measurement error. On the middle, each color corresponds to a single individual, with its theoretical functional response in solid line and its observations with measurement errors in dots. On the right, interaction stochasticity has been added to measurement errors: the solid line represents the mean functional response and the colored area represents the range of the distribution of the functional response.

## 2 Methods

The fundamental idea of the statistical approach presented hereafter is that data observed in an experiment, here the number of prey consumed by several isolated predators at different prey densities after a given time Δ, denoted by *R*, are the realizations of a stochastic process. In other words, we suppose that *R* follows a distribution which is not *a priori* chosen but that emerges from the process itself. This distribution constitutes the core of the statistical model used for inference. It converges to a Gaussian distribution with explicit mean and variance expressions [6] (note that it converges in law, which means that even though the number of preys eaten by a predator in a given time takes discrete values, it can be approximated by a Gaussian distribution which takes continuous values). Supposing that the distribution can be well approximated by this latter Gaussian distribution, we write a corresponding statistical model split into two main components: a deterministic part (the mean of the latter Gaussian distribution), and a stochastic part (including the variance of the latter Gaussian distribution).

In the following, we first present the statistical model including different sources of variation (measurement errors, between-predator variability, interaction stochasticity, Fig. 1). Second, we show that the two components of the statistical model naturally emerge from the convergence of the mechanistic model to an approximation of the distribution of *R* in general (Eq. 3), and then in a particular context and parameterization (Eqs. 4 and 5). Even though including a variance part to a statistical model is classical, either constant or dependent on the parameters of the model or the data structure, the form given to the variance is often an *a priori* or *adhoc* choice. Here on the contrary, the form given to the variance, and especially its dependency on the parameters, directly comes from the mechanistic stochastic model itself. The part of the variability emerging from the process originating the data (the interaction stochasticity, *i*.*e*. the mechanistic noise) is thus a source of information for parameter estimation.

### 2.1 A mechanistic stochastic model for estimating different variation sources in functional responses

#### Statistical model

We denote by *Y*_*ij*_ the random variable representing the number of preys consumed after a time Δ (considered as the unit of time, and determined by the experiment) by predator *i* at initial prey density or number *d*_*ij*_ (with *i* = 1, …, *N* and *j* = 1, …, *n*_*i*_) in a given experimental environment *e* (see Fig. 1). The statistical model that accounts for the different sources of variation under consideration is defined as the following non-linear heteroscedastic mixed-effects model:

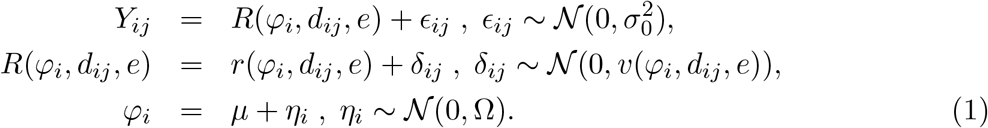

Here, the latent variable *R*, whose definition is detailed below, represents the interaction process between predators and prey. It is modeled as a deterministic function *r*, capturing the mean response, combined with a stochastic deviation *δ*_*ij*_, reflecting interaction stochasticity. Both *r* and the associated variance function *v* depend on the individual-specific traits *φ*_*i*_, which are modeled as random effects with population mean *µ* and variance Ω. Here, the individual-specific traits can be multidimensional, i.e. *ϕ*_*i*_ ∈ ℝ^*p*^, *µ* ∈ ℝ^*p*^ and Ω is a *p* × *p* matrix. Model (1) thus captures between-predator variability in behavior through the distribution of *φ*_*i*_, and incorporates process variance (via *δ*_*ij*_) and residual variance (via *ϵ*_*ij*_). Importantly, since the process variance *v*(*φ*_*i*_, *d*_*ij*_, *e*) results from a mechanistic approximation of the stochastic interaction model, it has a closed-form expression that depends explicitly on the model parameters and the form of the chosen functional response (see justification below). Assuming independence between these two sources of variations, the marginal distribution of *Y*_*ij*_ is Gaussian with mean *r*(*φ*_*i*_, *d*_*ij*_, *e*) and variance

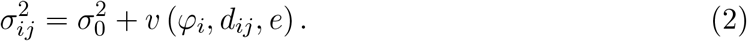

Since the variance of *Y*_*ij*_ depends on the individual predators traits *φ*_*i*_, prey density *d*_*ij*_, and environment *e*, the model is heteroscedastic by construction.

Note that even though *r* is referred to as the mean function, its connection to the population-level mean functional response for a given prey density is not direct. This is because the model is non-linear, and the individual-specific quantities *φ*_*i*_’s are random variables rather than fixed parameters. As a result, the expected functional response in the population is not simply obtained by evaluating *r* at *µ*. Instead, it involves integrating over the distribution of the random effects.

#### Theoretical justification

Our inferential model (Eq. 1) is derived from a mechanistic, microscopic (*i*.*e*. at the individual/predator level) and stochastic model describing how fast a single individual predator, with its peculiar set of traits, interacts with prey in a given environment during a foraging bout. Such a general model has been developed by [6] where they showed that given i) the density of prey *d*, ii) a set of predator traits *φ* (*e*.*g*. its foraging pattern, movement speed, handling time, probability of capture, etc.), iii) the distribution of the total times *T*_*k*_ taken by the predator to successfully eat the *k*^*th*^ prey during the foraging bout, and iv) an environment *e* potentially including prey traits variability, then the distribution of the functional response *R* (*φ, d, e*), *i*.*e*. the total number of prey consumed per unit of time during a foraging bout with a duration Δ, can be approximated, when Δ → ∞, by

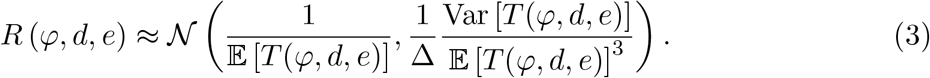

Approximation (3) is obtained under two main assumptions. First, that the times *T*_*k*_ are independent and have the same distribution as variable *T* (*φ, d, e*) in the above formula, which in particular implies that the variation of prey density *d* during the foraging bout is negligible (in other words, under the assumptions of renewal theory). In practice it means that the initial number of prey is assumed much larger than the number of eaten prey at the end of the foraging bout. Second, that the foraging bout duration Δ is large enough for a large number of successful interactions to have occurred. In that sense, Eq. (3) is a macroscopic approximation of the stochastic process underlying foraging. Such an approximation is necessary to make the model usable in the inference framework. Even though these assumptions might appear strong, especially regarding how experiments are performed, one should note that the functional responses classically fitted to data make, often implicitly, similar or even identical assumptions. For instance, Holling-type functional responses and their derived versions all implicitly suppose that prey are not depleted. Since they are all deterministic, they implicitly assume that the number of prey-predator interactions is very large, or equivalently that interactions are very fast processes. The functional responses accounting for prey depletion are also based on deterministic models, such as the RogersRoyama model, and the models derived by [31], thus with similar implicit assumption.

The model given by Eq. (3) can be considered as a general improvement for the inference of functional response because it makes it possible to derive both the mean *r* and the variance *v* of the functional response *R*, unlike in the case of deterministic functional response where only the mean is considered. More precisely, one can see the assumption that Δ → ∞ as a technical mean to derive the approximation. Δ is not effectively infinite, otherwise it would necessarily imply that the number of remaining prey at the end of the foraging bout is zero. In practice, this technical assumption makes it possible to find second-order approximation of the functional response: the first order approximation gives the mean *r*, while the second order gives the variance *v* (that is the reason why the variance is scaled to 1*/*Δ in Eq. (3)).

The functions *r* and *v* used to parameterize the inferential model (Eq. 1) are precisely obtained as the mean and variance of the distribution specified in (3). Eq. (3) thus helps to disentangle different sources of variation expressed in the inferential model (Eq. 1). As Eq. (3) was derived at the individual level, it allows to quantify both the expectation of the functional response *r* and its random fluctuations due to the foraging process itself *v*, considering the possibility that each individual can have its own set of traits (thus accounting for between-predator variability). Estimating *v*, the variation due to the foraging process itself, is made possible because the size of the random fluctuations of the functional response depends on the individual predator traits *φ*, under the condition that collecting data from the same single individual is possible.

#### Why Gaussian distributions?

In the statistical model in Eq. (1), the sources of variation are all modeled as Gaussian distributions. There are two distinct reasons why this is so depending on the source of variations. On the one hand, it can correspond to *a priori* assumptions like for the residual error *ϵ*_*ij*_ and the between-predator variability *η*_*i*_. The Gaussian assumption is very common and typical for such variables in the ecology and statistics literature, albeit not mandatory. On the other hand, the Gaussian assumption can also arise as an emergent property. This is the case for the variability due to interaction stochasticity *v*: the Gaussian distribution naturally emerges from the stochastic process model that underlies the functional response and its approximation (see the section ‘Theoretical justification’). A Gaussian distribution as an approximation of the distribution of the functional response makes sense for two reasons. First because the functional response is defined as a rate, a number of prey eaten in a given time, and need not take only discrete values. Second, according to the Central Limit Theorem, it is expected that the Gaussian distribution is generally a good approximation as long as the sample size is large (which corresponds here to the assumptions that the foraging duration Δ is large).

Yet, the statistical framework we describe here is general enough to be adapted to other options depending on the experimenters’ knowledge and objectives. Eq. (1) can indeed be modified accordingly to include other distributions, such as in [31] where individual traits are randomly drawn from a uniform distribution. In addition, the Stochastic Gradient Descent procedure is general and robust enough for estimation and optimization in many situations, including distributions that do not belong to the exponential family. In practice, non-Gaussian distributions can be *a priori* chosen for the residual error *ϵ*_*ij*_ and the between-predator individual variability *η*_*i*_ in Eq. (1).

The distribution followed by the functional response *R* itself can also be *a priori* chosen, including discrete distributions such as the Poisson distribution. However, this would necessarily imply an *a priori* form for the variance of the functional response with the need for an *a posteriori* mechanistic justification. Our statistical framework can also be used with other distributions that would emerge from the approximation of another stochastic process, given there is an explicit form for this distribution (note that this is not necessarily the case, as for instance when spatial foraging and depletion are both considered [4]).

#### Example

In order to illustrate the derivation of a statistical model from Eq. (3) and its use for performing inferences from a dataset, we will consider in the numerical illustrations of the paper a specific simplified context which is yet classically used in ecology. We suppose that a single predator forages in a constant environment *e* containing identical prey, with prey density *d* fixed by the experiment design. For the sake of clarity, since there is only one predator we omit the index *i* in this paragraph. The predator traits set is characterized by its searching rate *λ* and the time it takes for handling a prey when captured *h, i*.*e. φ* = (*λ, h*), and we suppose that its foraging pattern is simply that it goes directly to the closest available prey. [6] showed that under these assumptions (similar to the Holling type II model [16], which is deterministic) then an approximation of the stochastic functional response is given by a Gaussian distribution with mean

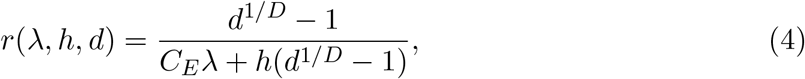

and variance

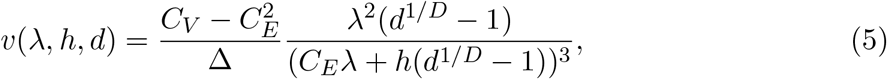

where *C*_*E*_ and *C*_*V*_ are known constants that only depend on the dimension *D* of the environment (see [6]). Note in particular that when Δ → ∞, the variance vanishes as *v* → 0 in Eq. (5), and one recovers Holling-type deterministic functional response as the mean *r* in Eq. (4). Also note that the −1 in the numerator and denominator comes from the approximation under the assumption that the number of interactions should be large enough, thus impliying that the number of initial prey *d* ≫ 1. Δ is the time scale of the experiment which is fixed to Δ = 1 in the following for simulated datasets without loss of generality, so that *r* is to be expressed as the number of prey ingested by the predator per unit of time Δ. In the case of the real dataset, it was fixed to Δ = 2 min as it was the duration of an experimental sequence. In this model, 1*/λ* represents the speed of prey capture per unit area and 1*/h* represents the asymptotic number of prey a predator can ingest when the prey density tends to infinity. For simplicity we did not consider environmental variability in this example, thus assuming that all measurements are made under the same experimental conditions. The environment *e* is therefore omitted from the notations above.

### 2.2 Statistical Tools for Mixed-Effects Models with Heteroscedasticity

A fundamental point for inference is that, in mixed-effects models—particularly in the one defined by Equation (1)—the random effects *φ*_*i*_ are not model parameters. These quantities, also sometimes called ‘individual parameters’, are random variables that cannot be directly observed. Therefore, the vector of parameters to be estimated is *θ* = (*µ*, Ω, 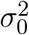). Note that *µ* is a vector of *p* parameters and Ω is a *p* × *p* matrix. Estimating these parameters makes it possible to characterise and quantify the average behaviour in the population, the between-predator variability, and the residual variance. Inference therefore requires dedicated statistical tools, not only because of the presence of latent variables but also because of the model’s non-linearity. Indeed, the individual-specific random effects *φ*_*i*_ enter the model non-linearly, both in the mean function *r* and in the variance function *v*. As a result, standard estimation procedures based on explicit likelihoods or simple sufficient statistics are no longer tractable. Likelihood-based inference must instead integrate over the distribution of the latent random effects, which introduces additional computational and methodological challenges. Indeed, the likelihood is defined as the joint distribution of the observed data; in our case the observed data are the *y*_*i*_, for *i* = 1, …, *N*, where *y*_*i*_ gathers all the observations of predator *i*, i.e. 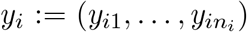. Note that the notation *y*_*i*_ refers to the realization of the underlying random variable which was denoted by *Y*_*ij*_ in model (4). However, we don’t have access to the marginal distribution of the *y*_*i*_, but only to the conditional distribution of *y*_*i*_ given the individual parameters *φ*_*i*_ and ii) the marginal distribution of the individual parameters *φ*_*i*_, the product of which being equal to the joint distribution of *y*_*i*_ and *φ*_*i*_. Using Bayes theorem, the marginal distribution of the *y*_*i*_ is given by the integral of the joint distribution, over the random variables *φ*_*i*_, without an explicit solution here. Moreover, although the theoretical framework for mixed-effects models is well established (see, for instance, [29] and [21]), no generic software currently supports mixed-effects models that combine inter-individual variability with heteroscedasticity, especially when the latter arises from a non-linear variance structure derived from a mechanistic stochastic process. In the following, we present statistical tools specifically designed for parameter estimation and model comparison in functional response models that incorporate multiple sources of variation, with the aim of making them accessible and practical for experimenters.

#### 2.2.1 Algorithms for parameter estimation

##### A brief review of the state of the art

Several methods have been developed to tackle parameter estimation in non-linear mixed-effects models. However, two particularly powerful and flexible likelihood-based strategies stand out for our context: the Stochastic Approximation EM (SAEM) algorithm [13, 20] and Stochastic Gradient Descent (SGD) [1]. SAEM approximates the complete-data likelihood through stochastic updates, while SGD seeks to drive unbiased estimates of the gradient of the log-likelihood toward zero, and is often easier to implement. Importantly, an additional advantage of SGD is that, beyond parameter estimation, it naturally provides an estimate of the Fisher information matrix as a by-product of the optimization process. This output is valuable for uncertainty quantification. Detailed descriptions of both algorithms are provided in Appendix A.

SAEM is implemented in several software packages, including the `saemix` package for R [9]. A key limitation of current implementations is that they only support heteroscedastic models where the variance depends on the mean through simple parametric forms—typically the proportional error model 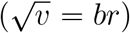 or the combined error model 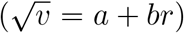, with constants *a* and *b* to be estimated. More general forms of heteroscedasticity, such as those arising from mechanistic stochastic processes, are not covered. In these cases, the model no longer belongs to the exponential family, and the required updates at each iteration of the algorithm do not admit closed-form expressions (see [20]). This significantly increases the implementation complexity and computational cost of SAEM (see Appendix A). In contrast, to our knowledge, there are no dedicated R or Python packages for SGD in mixed-effects models. We therefore developed a tailored code to enable statistical inference in functional-response models under this framework. Appendix A details the implementation, and the full source code used for numerical experiments and data analysis is available on GitHub at https://github.com/baeyc/functional_responses.

##### Practical considerations for SGD

Applying Stochastic Gradient Descent in non-linear mixed-effects models involves several practical considerations to ensure reliable and efficient convergence. As an iterative method, SGD progressively refines parameter estimates over successive updates, starting from initial values. The final estimate corresponds to the last iteration, under the assumption that convergence has been achieved. The performance of SGD depends on a careful tuning of algorithmic parameters and diagnostic monitoring throughout the estimation process. A first critical aspect is initialization: because SGD tends to converge to a local rather than a global optimum, initial values should be chosen carefully, ideally informed by domain knowledge, to guide the algorithm toward plausible and interpretable solutions. Another key factor is the choice of the learning rate (denoted by *γ*_*k*_ in Appendix A), which controls the magnitude of parameter updates. A large initial learning rate can promote exploration of the parameter space, helping to avoid poor local optima, while a decreasing learning rate allows for finer adjustments as the algorithm converges. Designing an appropriate learning rate evolution therefore requires balancing between exploration and stability; (detailed guidelines can be found in [1, 25]). In addition, convergence diagnostics play an important role in practice. Plotting the evolution of the estimated parameters or the log-likelihood across iterations can help assess whether the algorithm is stabilizing appropriately. Signs of instability may indicate the need to revise the initialization or adjust the learning rate. Figure 2 illustrates such behavior in a representative example. In order to be confident enough in the inferences made with data from the model, the algorithm should properly converge and estimate values in accordance with data, and the robustness and consistency of the estimates should be assessed. These are necessary conditions before moving forward to comparing different models, selecting an appropriate noise structure, or determining whether a parameter should include inter-individual variability.

**Figure 2:**
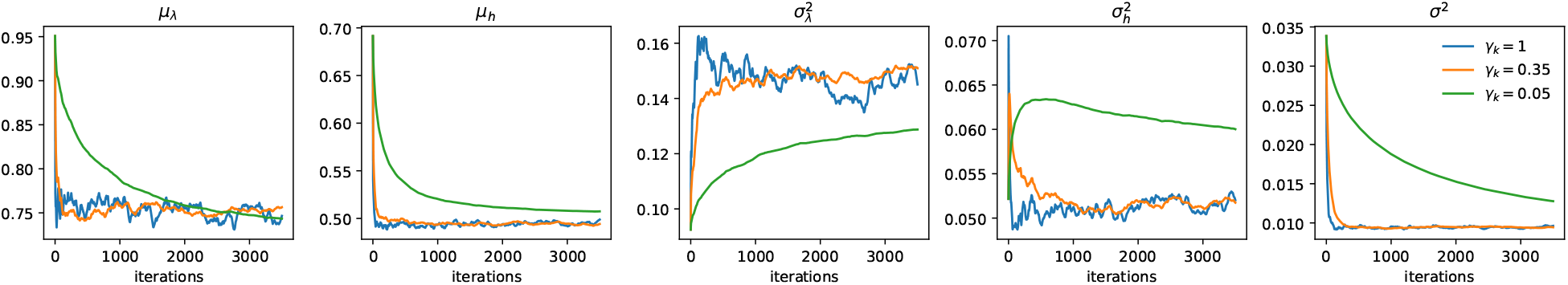
Effect of the learning rate on the convergence of the Stochastic Gradient Descent algorithm for estimating the parameters of the model. When the learning rate *γ*_*k*_ is too small (in green), the trajectory is smooth but the convergence is too slow. As the step size increases, the algorithm achieves a faster convergence but the trajectory exhibits higher variability.

Note that even though individual effects *φ*_*i*_ are not parameters to be estimated, it is still possible to predict their values. One approach proceeds as follows. Since the algorithms used to estimate *θ*—described above and detailed in the Appendix A—rely on simulating candidate values for these individual effects, predictions for the *φ*_*i*_’s can be obtained as a by-product of these procedures. Specifically, they can be retrieved from the simulated values generated during the final iterations of the algorithm, denoted by 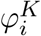 in Appendix A.

#### 2.2.2 Model comparisons

In the context of nonlinear mixed-effects models, information criteria such as the Bayesian Information Criterion (*BIC*) provide a valuable and practical framework for model selection. While alternative approaches and more classical statistical tests (e.g., likelihood ratio tests) exist, these often involve significant analytical and computational challenges due to the models’ hierarchical structure, where individual-level parameters are latent random effects, and their intrinsic non-linearity. Such complexity leads to test statistics that may follow non-standard distributions, making their evaluation both analytically intractable and numerically expensive, especially when determining appropriate covariance structures for the random effects [2].

Moreover, the *BIC* offers the advantage of enabling comparison between models making assumptions on any source of variability, be it in the mean structure, variance components, or random effects, whereas many classical tests or alternative tools tend to be tailored to specific hypotheses or focused on particular model components. For these reasons, we focus here on information criteria as a simpler, more intuitive, and computationally efficient tool for model comparison, without the need for costly *p*-value estimation.

##### Illustration of models comparison: is interaction stochasticity significant?

We illustrate model comparison and selection with our framework by tackling the following issue: is the variability due to interaction stochasticity a significant part of the total variability observed in data? To answer this question we compared candidate models that differ by including or excluding the stochastic component derived from the mechanistic formulation

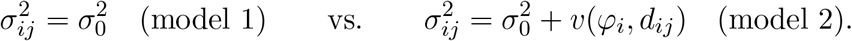

and compared which one fits best to data using the *BIC* criterion

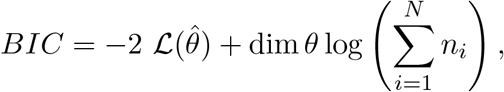

where 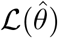 denotes the log-likelihood evaluated at the estimated parameters, dim *θ* is the number of estimated parameters, and 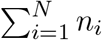 is the total number of observations. Comparing models based on their *BIC* values by selecting the one with the smallest criterion, provides a straightforward and unified decision rule, regardless of the nature of the variability components. The main technical challenge here lies in the evaluation of the log-likelihood, which is not available in closed form due to the presence of latent random effects and the complexity of the variance structure. To overcome this, we rely on Monte Carlo approximations of the likelihood, as detailed in Appendix A.2.

Comparing and selecting between the two models with and without interaction stochasticity is supposedly possible as the individual parameters *λ*_*i*_ and *h*_*i*_ appears both in the mean and the variance of the Gaussian distribution approximating the functional response in Eq. (1). As a consequence, the mean and variance of the number of eaten prey are not independent, whereas the residual noise is assumed to be independent of the functional response and hence of its parameters. This should thus add some constrains in parameter estimation that should make it possible to distinguish between the two models.

### 2.3 Numerical experiments

#### Objective

We conducted a simulation study in order to illustrate how experimental design and data-generating conditions affect the quality of parameter estimation and model selection in heteroscedastic mixed-effects models, particularly with respect to the identification and estimation of different sources of variability, and indirectly to provide guidelines in designing experiments. To facilitate analysis and interpretation, we considered a two-dimensional model with random parameters that do not depend on covariates. Under a fixed measurement budget, we examined how the trade-off between the number of individuals and the number of repeated measurements per individual influences estimation bias, estimator variance, and model selection based on the Bayesian Information Criterion (BIC). We also investigated how the magnitude of variability—both between and within individuals—affects these performance metrics.

#### Model

We generated data following the statistical model defined in Eqs. (1)-(2) where the observations variance is splitted into a mechanistic term *v* and a residual term 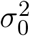. *r* is given by Eq. (4) and *v* by Eq. (5) with environment dimension *D* = 2. Accordingly to the assumption of the statistical model (1), the two individual parameters *λ*_*i*_ and *h*_*i*_ were randomly drawn for each individual *i*, from a Gaussian distributions with fixed mean *µ*_*λ*_, *µ*_*h*_, and fixed variance 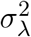 and 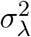, respectively. In addition, we introduced a covariance *σ*_*λ,h*_ between the two traits in order to take into account that handling times and searching rate might not necessarily be independent, which is expected if those traits both depend on the same covariates such as individual’s size, weight or experience (*e*.*g*. [34].) A log-transformation was applied to the two behavioral parameters to account for their positiveness. More precisely, we assumed the following:

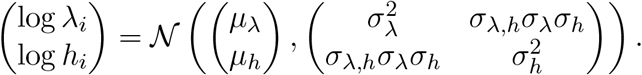

The model is associated with typical curve shapes that vary according to the parameter values. Consequently, depending on the densities at which observations are collected, they are likely to provide information on all or part of the curve, resulting in better or poorer quality estimates. Two example datasets associated with different parameter sets are given in Figure 3, with 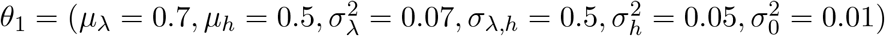 in the first case and with 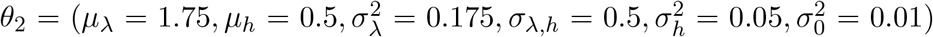 in the second case. The asymptotic number of prey a predator can ingest is denoted by 1*/h* and is identical in both settings. The difference between the two parameter sets lies in the value of the behavioral parameter *λ*, which influences the speed of prey consumption. For a prey density that varies between 1 and 150, we can observe the whole functional response trajectory with parameter set *θ*_1_, while the asymptotic regime is not reached with the second parameter set *θ*_2_.

**Figure 3:**
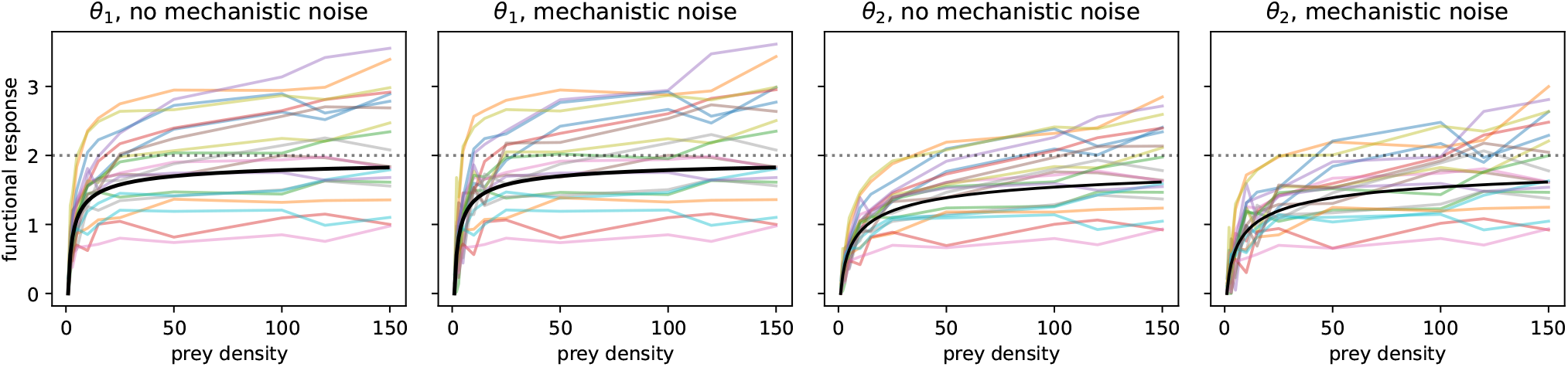
Example of simulated datasets using *θ*_1_ and *θ*_2_, with and without interaction stochasticity. The dotted line represents 1*/h*, the asymptotic number of prey a predator can eat. The black curve represents the mean functional response in the population. The different curves represent functional responses for different individual predators, each with their own individual traits *λ* and *h*.

The generation of simulated data thus requires two types of inputs: i) a set of parameter values from which the functional response can be derived, and ii) the observation scheme, *i*.*e*. the prey densities at which the data are to be observed. Note that, given that our objective here is to provide a proof-of-concept that our statistical framework can disentangle and estimate different sources of variation, we do not compare different functional responses, but different variance structures for the same functional response. In this sense, the *a priori* chosen functional response can be considered as a third extra type of inputs.

#### Experimental settings

In all cases below, we evaluate the quality of the estimates and the possibility of choosing the right model when comparing the model with or without the variability from interaction stochasticity *v*(*φ*_*i*_, *d*_*ij*_) by means of the BIC (*i*.*e*. we compare a model with variance 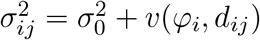 to a model with variance 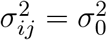).

1. We first considered several configurations for the number of individuals *N* and the number of measurements per individual *n* (we therefore assumed the same number of measurements for each individual). For each parameter set, *θ*_1_ and *θ*_2_ above, and for a fixed total number of measurements of approximately *nN* = 500, we let the number of individuals and the number of measurements vary as (*N, n*) ∈ {(10, 50), (20, 25), (30, 17), (40, 12), (50, 10), (60, 8), (70, 7), (80, 6), (90, 5)}. The prey densities at which the data are to be observed were randomly generated between 1 and 150, for each value of *n*. The same set of generated values was then used throughout the numerical experiments and was thus considered as fixed.
2. We then investigated the influence of the signal-to-noise ratio through two complementary experiments. First, we varied the coefficient of variation of the random effects in {25%, 50%, 75%}, thereby increasing the relative dispersion of individual parameters around the population mean. Second, we modulated the measurement noise by adjusting the residual standard deviation *σ*_0_, expressed as a proportion of the asymptotic response, using values of 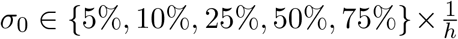. These experiments are conducted for parameter set *θ*_1_ only, assuming zero covariance among the random effects.

#### Evaluation criteria

We assess the estimation accuracy by measuring the bias and the variance of the estimates obtained on the simulated datasets, which can then be combined in the mean squared error (MSE). For that purpose, for a given parameter set *θ* and for each tested sample size, we generated *K* = 1000 datasets and computed the corresponding estimate 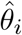 for each dataset. We then computed the aforementioned criteria as:

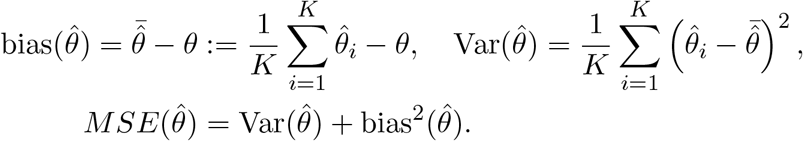

In order to compare the estimates of parameters that can be defined on different scales, one can use the relative root MSE, which is defined as 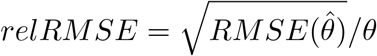. Model selection is assessed by counting the number of times, out of *K* = 1000 repetitions, that the correct model is actually found, *i*.*e*. associated with a smaller BIC value.

### 2.4 Real dataset

We ran our inferential model on a real dataset from [34] (see Figure 4, data available in [19]). The number of invertebrate prey (*Artemia salina*) ingested by an isolated predator fish (*Heterandria formosa*) in two minutes was measured by direct visual observation at 14 different initial prey densities (range: 10-2000 prey per 2L). Forty-nine different predator individuals were used for the experiment. The number of prey ingested was measured for the 14 initial prey densities for all individual predators (*i*.*e*. a given individual predator was measured 14 times independently, which gives information about the variation due to interaction stochasticity). Two feeding trials were performed per day and a trial series lasted 7 days. The duration of one feeding trial being Δ = 2 minutes, it gives the time dimension of the estimated parameters as they are scaled to Δ in Eq. (3).

**Figure 4:**
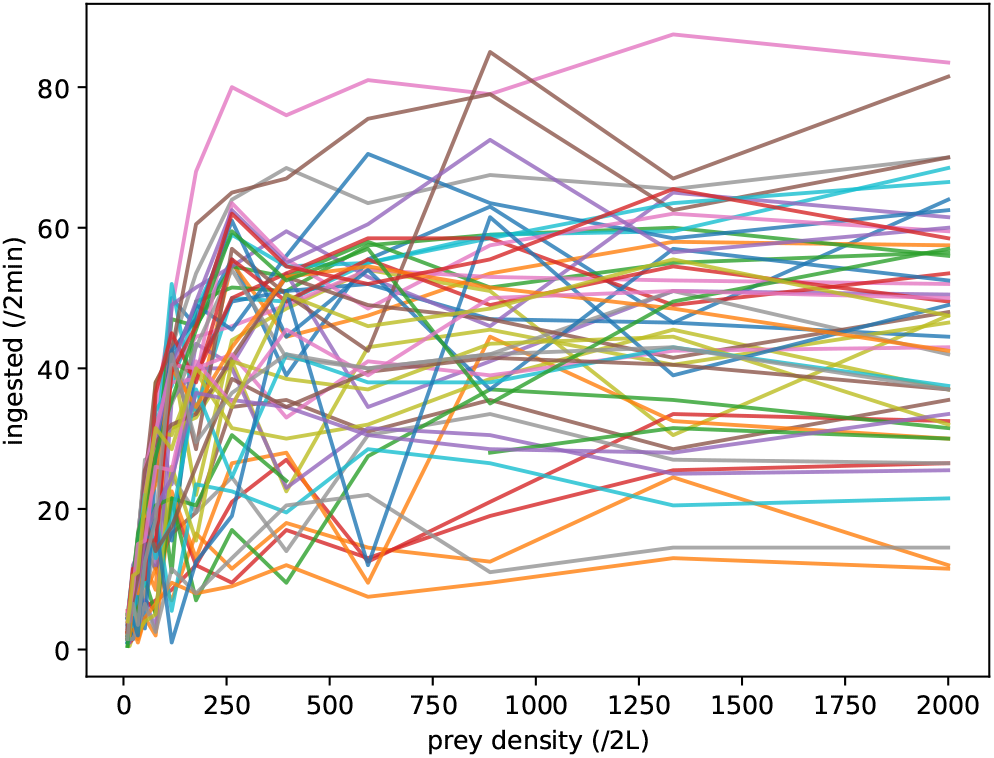
Number of ingested *A. salina* by a single predator *H. formosa* during a time window of 2 minutes in a 2L aquarium, at 14 different food densities (see [34]). Each color corresponds to the measurements made on the same single predator at different prey densities.

#### Estimation and validation procedures

We fitted the 3-dimensional (*D* = 3, Eqs. 4 and 5) functional response model to the *A. salina* (prey) vs. *H. formosa* (predator) data, with *λ* and *h* as correlated random effects and considering a mechanistic term in the residual variance (model 2). As the SGD algorithm can converge to a local optimum, we ran ten repetitions of the algorithm with random initialization. As results were very similar for all runs (see Figs. A.1 and A.2), we averaged the estimates from the 10 runs.

To assess the ability of the model to capture the variability of the population, several diagnostic plots were performed. First, individual predictions were compared to observed data (see Fig. 12 for six randomly selected individuals and Fig. A.3 for all individuals), where the individual predictions were obtained using the predicted individual parameters 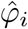 and the mean functional response *r*, i.e. the individual predictions for individual *i* at prey density *j* is 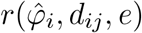 (see Eq. 4). Estimations of the individual parameters were obtained as a by-product of the SGD algorithm, using the samples 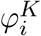 obtained at the last iteration of the algorithm. To account for the interaction stochasticity, we simulated 100 realizations of the stochastic model for each individual. Second, we used the Visual Predictive Check (VPC) [42] to assess the ability of the model to correctly reproduce the observed data (Fig. 13). In a VPC, the theoretical quantiles obtained from the model are compared to the empirical quantiles obtained from the observed data. More precisely, the procedure is as follows: i) we generated 5000 simulated datasets of 49 individuals (i.e. the same sample size as the observed data) under the model with its estimated parameters; for each simulated dataset and each prey density, we computed the 5%, 50% and 95% quantiles of the number of ingested prey over the 49 individuals of the population; and iii) we computed the 5%, 50% and 95% quantiles of the quantities obtained at the previous stage over the 5000 repetitions.

#### One-step *vs*. two-steps estimation approaches

The sources of variation are jointly considered in the statistical model in Eq. (1). It means that the parameters involved in the between-predator variability, the residual variance, the effect of the treatment (the initial density of prey), and the variability from interaction stochasticity are estimated in a single step jointly using all sources of variation from data. This one-step approach is to be compared to two-steps approaches often used in the literature in order to estimate the individual parameters between predator *e*.*g*. [34]. In such a two-steps procedure, functional responses are first fitted for each individual separately and then merged to estimate parameters at the population level. Such a two-steps approach has several limitations due to the fact that statistical errors tend to propagate, hence decreasing the accuracy as well as increasing biases of estimations [36]. In order to illustrate that specific point, we estimated parameters of our model using the one-step vs. two-steps approaches on [34]’s dataset and compared the results.

## 3 Results

### 3.1 Estimation accuracy

#### Experimental designs comparison

Results are given in Figs. 5 and 6 for both parameter sets *θ*_1_ and *θ*_2_. Different behaviors can be observed for the relative MSE. In general, the relative MSE decreases as the number of individuals increases. However, for some parameters we observed a first decreasing phase followed by a rising phase. We recall that the total number of measurements is maintained approximately constant around 500, so that when the number of individuals increases, the number of measurements per individual decreases. The latter behaviour could thus be explained by the fact that in those configurations, the number of measurements per individual is too small to correctly estimate the corresponding parameter. This is the case for example for parameters *µ*_*λ*_ and *µ*_*h*_ when the data were simulated using *θ*_2_, starting from *N* = 50, *n* = 10, and to a lesser extent, for parameter *µ*_*λ*_ when the data were simulated using *θ*_1_, starting from *N* = 70, *n* = 7. In these settings, we observe that as *µ*_*λ*_ is underestimated, *µ*_*h*_ is overestimated, suggesting that compensation can occur when the number of measurements per individual is too small. This is particularly true when the observation scheme does not cover the asymptotic regime, as this is the case with *θ*_2_.

**Figure 5:**
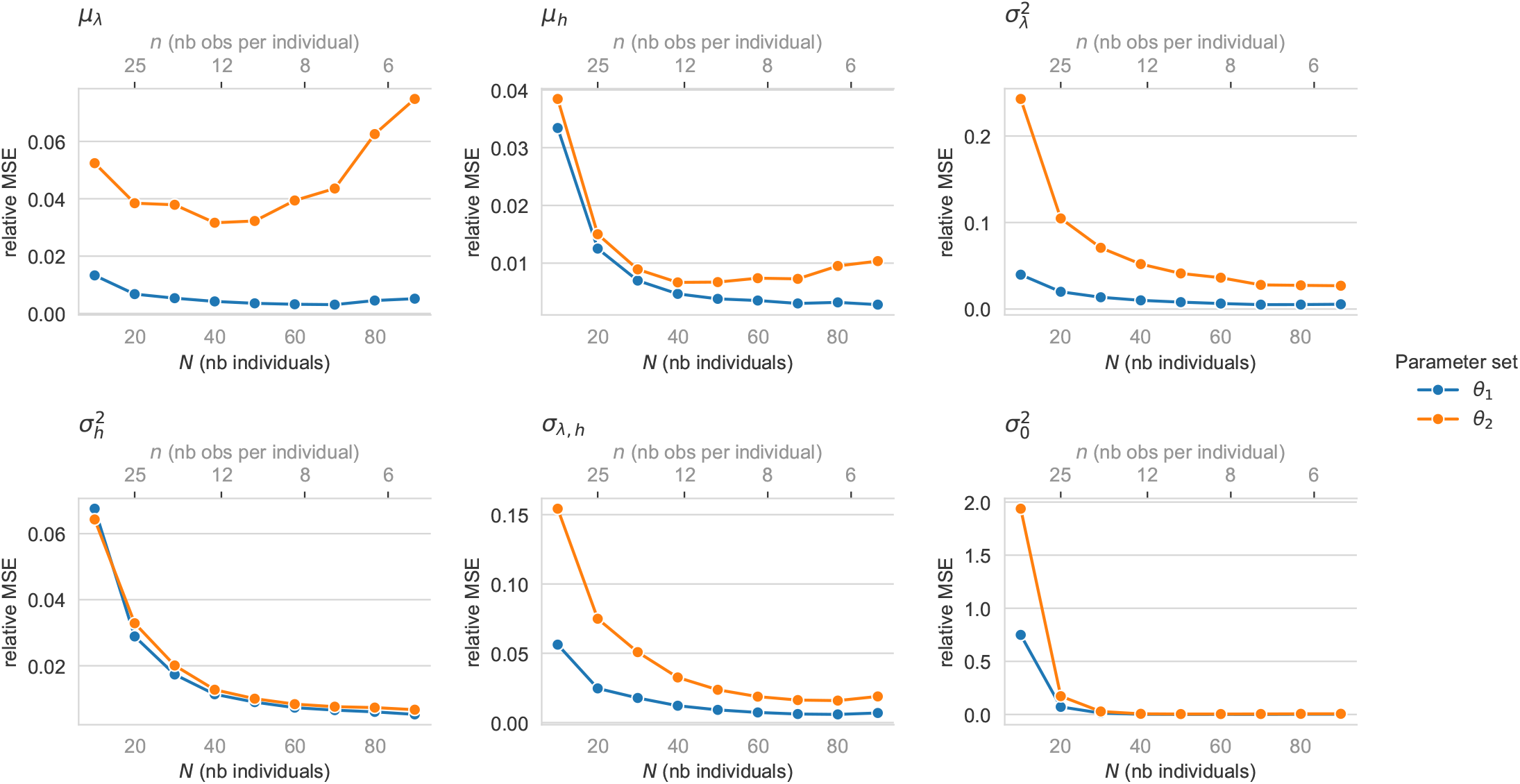
Relative Mean Squared Error (MSE) of each parameter as a function of the number of individuals in the population, for both parameter sets *θ*_1_ and *θ*_2_, computed over *K* = 1000 repetitions.

**Figure 6:**
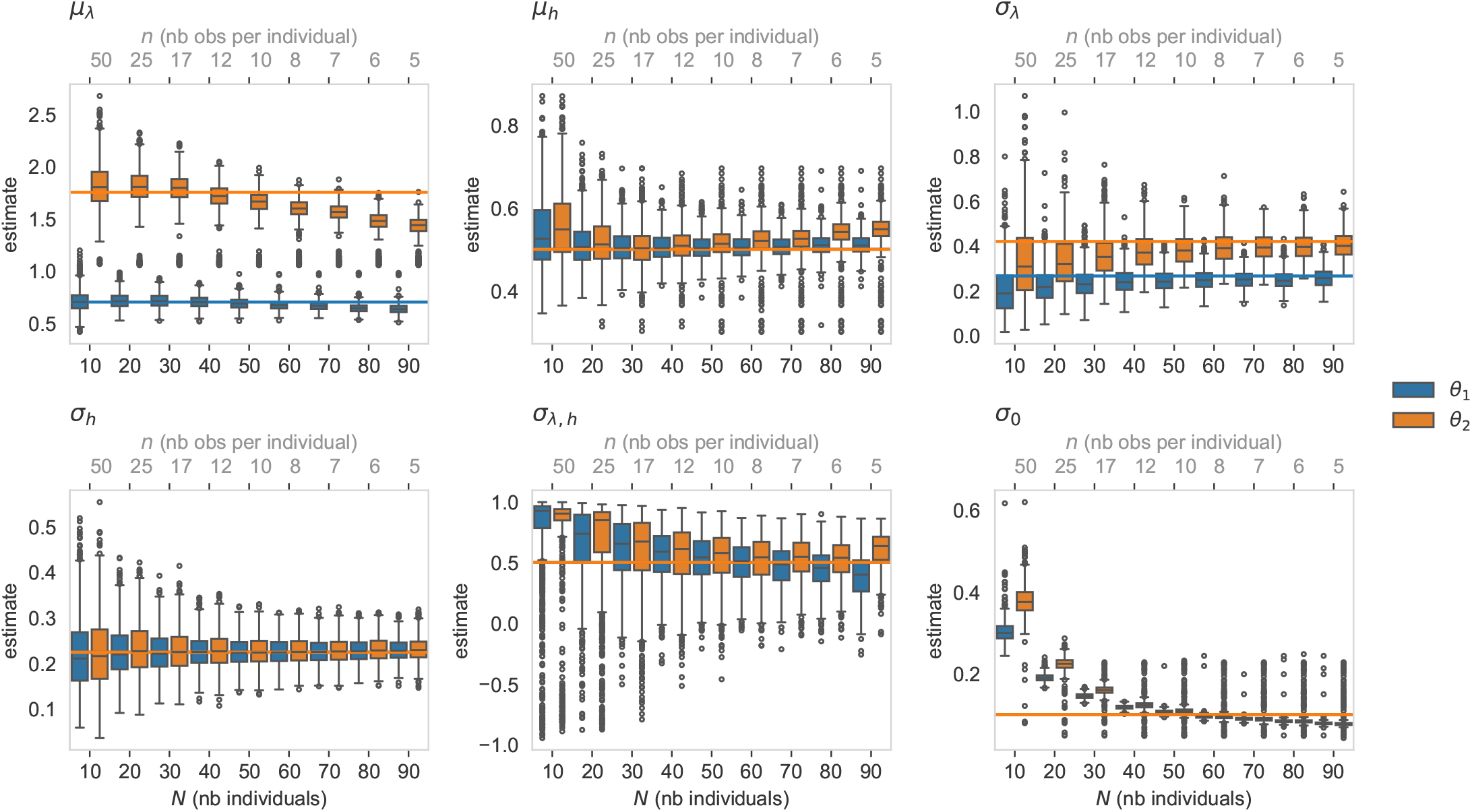
Boxplot of the estimates obtained on *K* = 1000 repetitions, as a function of the number of individuals in the population (principal x-axis) and of the number of observations per individual (secondary x-axis), for the two parameter sets *θ*_1_ (orange) and *θ*_2_ (blue). The horizontal line shows the true value of each parameter under each parameter set.

#### Effect of the signal-to-noise ratio

Results are given in Figs. 7, 8, 9 and 10. When the residual variance increases (Figures 9 and 10), all model parameters tend to be estimated with larger bias. The estimation uncertainty as measured by the standard deviation of the estimators, also increases with higher residual variance. A similar effect is observed with increasing random effect variance (Figures 7, 8). This suggests that reducing the measurement errors can directly improve the quality and precision of estimates and inference, a gain to be balanced by the associated experimental costs. An alternative could also be to better control the inter-individual variability which might be difficult to achieve as it is often inherent to biological systems, and which might even not be desirable if evaluating the role of inter-individual variability is one of the aim of the experiment.

**Figure 7:**
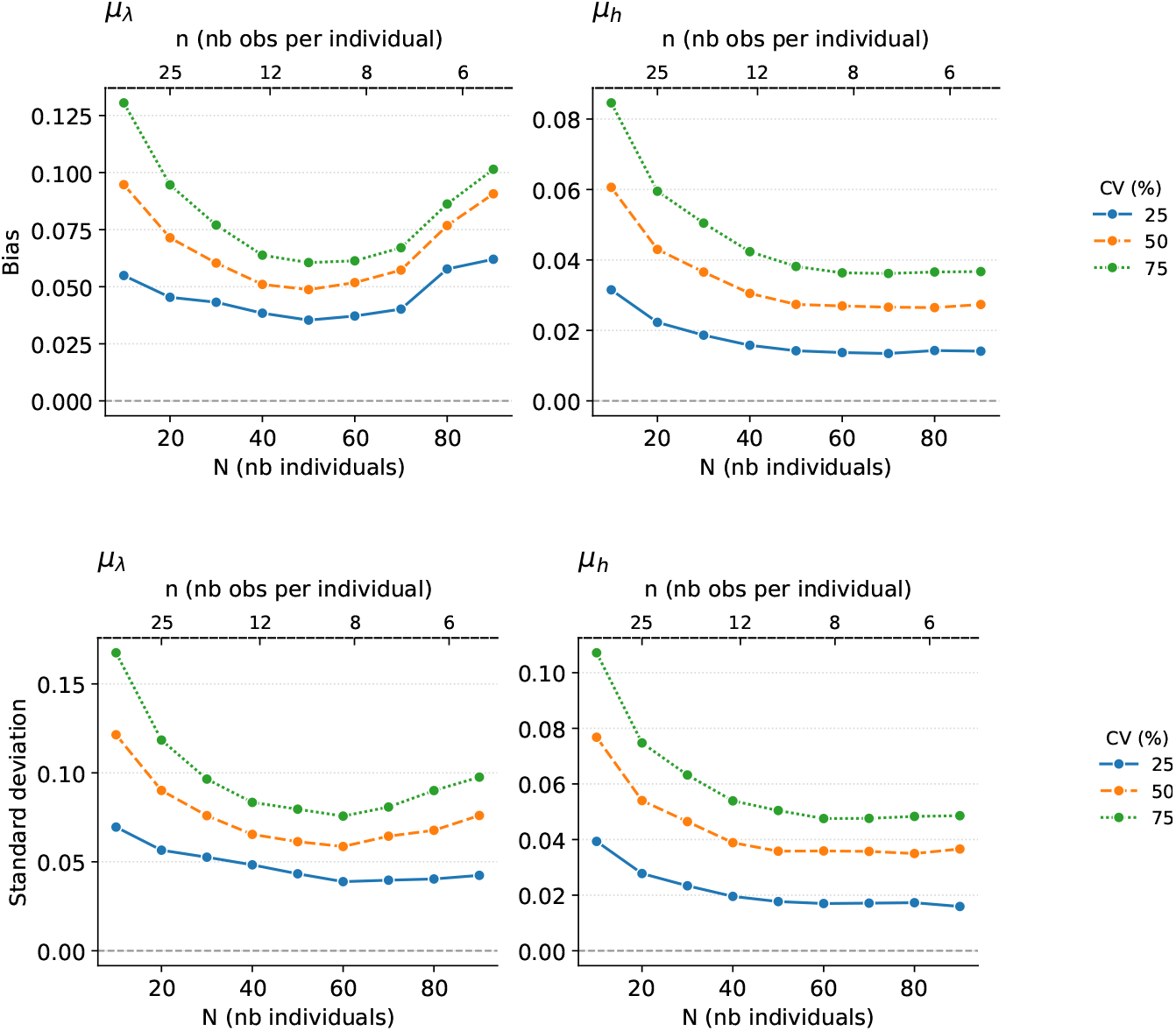
(top) Bias (in absolute value) of each mean parameter as a function of the sample structure, *i*.*e*. the number of individuals in the population (principal x-axis) and the number of observations per individual (secondary x-axis), for different coefficients of variation for the random effects and the parameter set *θ*_1_, computed over *K* = 1000 repetitions. (bottom) Standard deviation of each mean parameter as a function of the sample structure, *i*.*e*. the number of individuals in the population (principal x-axis) and the number of observations per individual (secondary x-axis), for different coefficients of variation for the random effects and the parameter set *θ*_1_, computed over *K* = 1000 repetitions.

**Figure 8:**
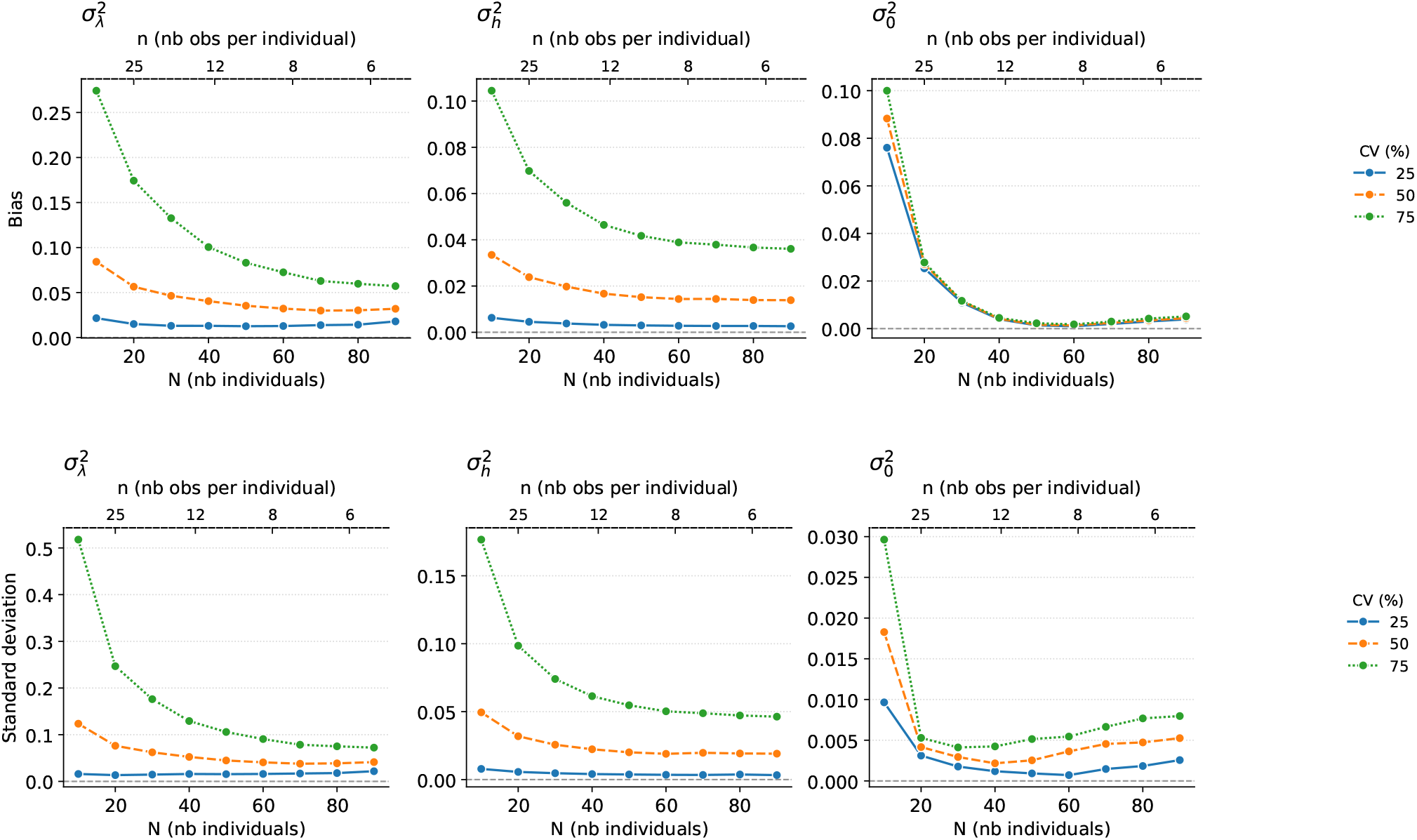
(top) Bias (in absolute value) of each variance parameter as a function of the sample structure, *i*.*e*. the number of individuals in the population (principal x-axis) and the number of observations per individual (secondary x-axis), for different coefficients of variation for the random effects and the parameter set *θ*_1_, computed over *K* = 1000 repetitions. (bottom) Standard deviation of each variance parameter as a function of the sample structure, *i*.*e*. the number of individuals in the population (principal x-axis) and the number of observations per individual (secondary x-axis), for different coefficients of variation for the random effects and the parameter set *θ*_1_, computed over *K* = 1000 repetitions.

**Figure 9:**
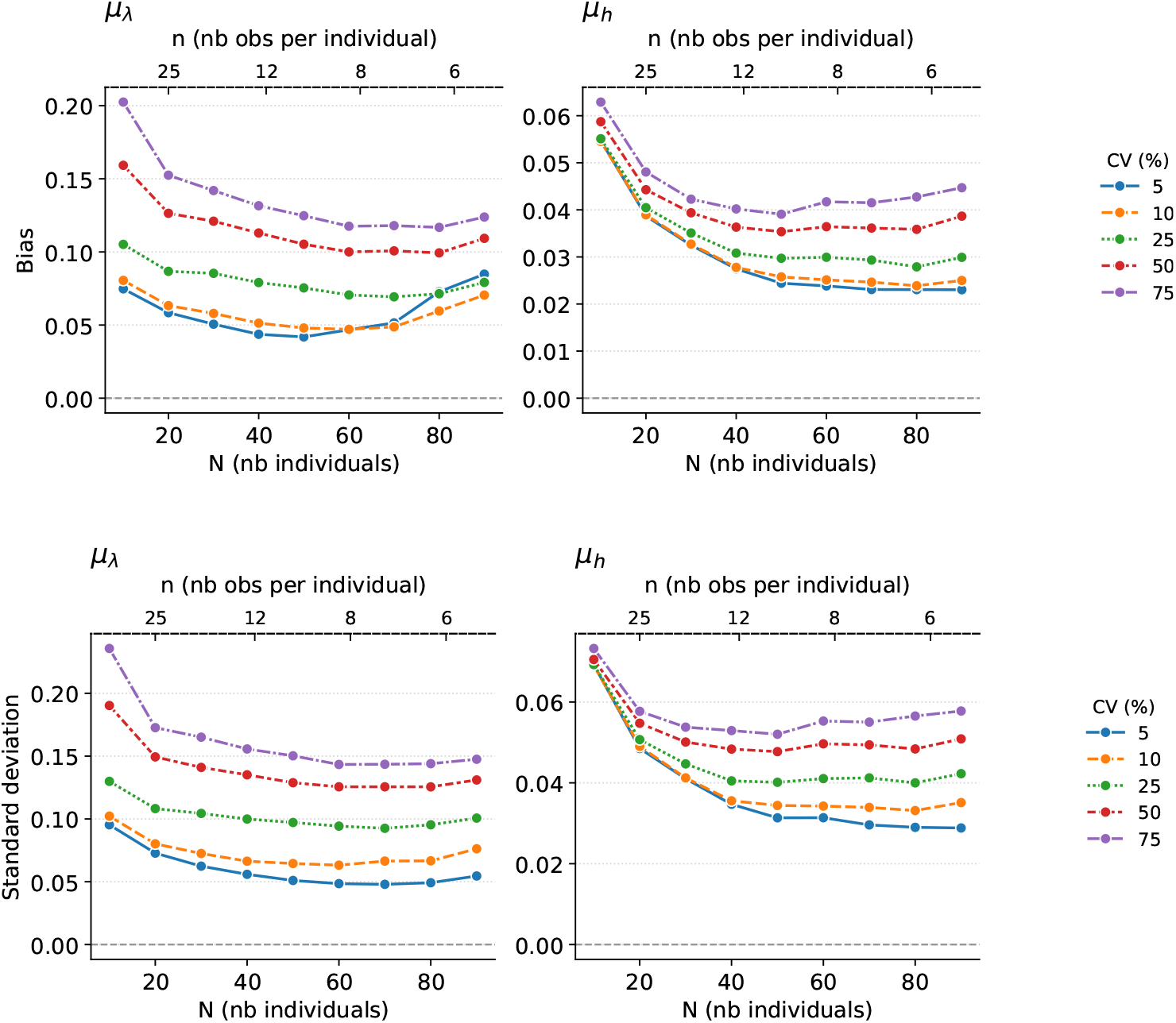
(top) Bias (in absolute value) of each mean parameter as a function of the sample structure, *i*.*e*. the number of individuals in the population (principal x-axis) and the number of observations per individual (secondary x-axis), for different coefficients of variation for the residual variability and the parameter set *θ*_1_, computed over *K* = 1000 repetitions. Only values with biases greater than 1 are shown (for reasons of scale). (bottom) Standard deviation of each mean parameter as a function of the sample structure, *i*.*e*. the number of individuals in the population (principal x-axis) and the number of observations per individual (secondary x-axis), for different coefficients of variation for the residual variability and the parameter set *θ*_1_, computed over *K* = 1000 repetitions. Only values with standard deviations greater than 1 are shown (for reasons of scale).

**Figure 10:**
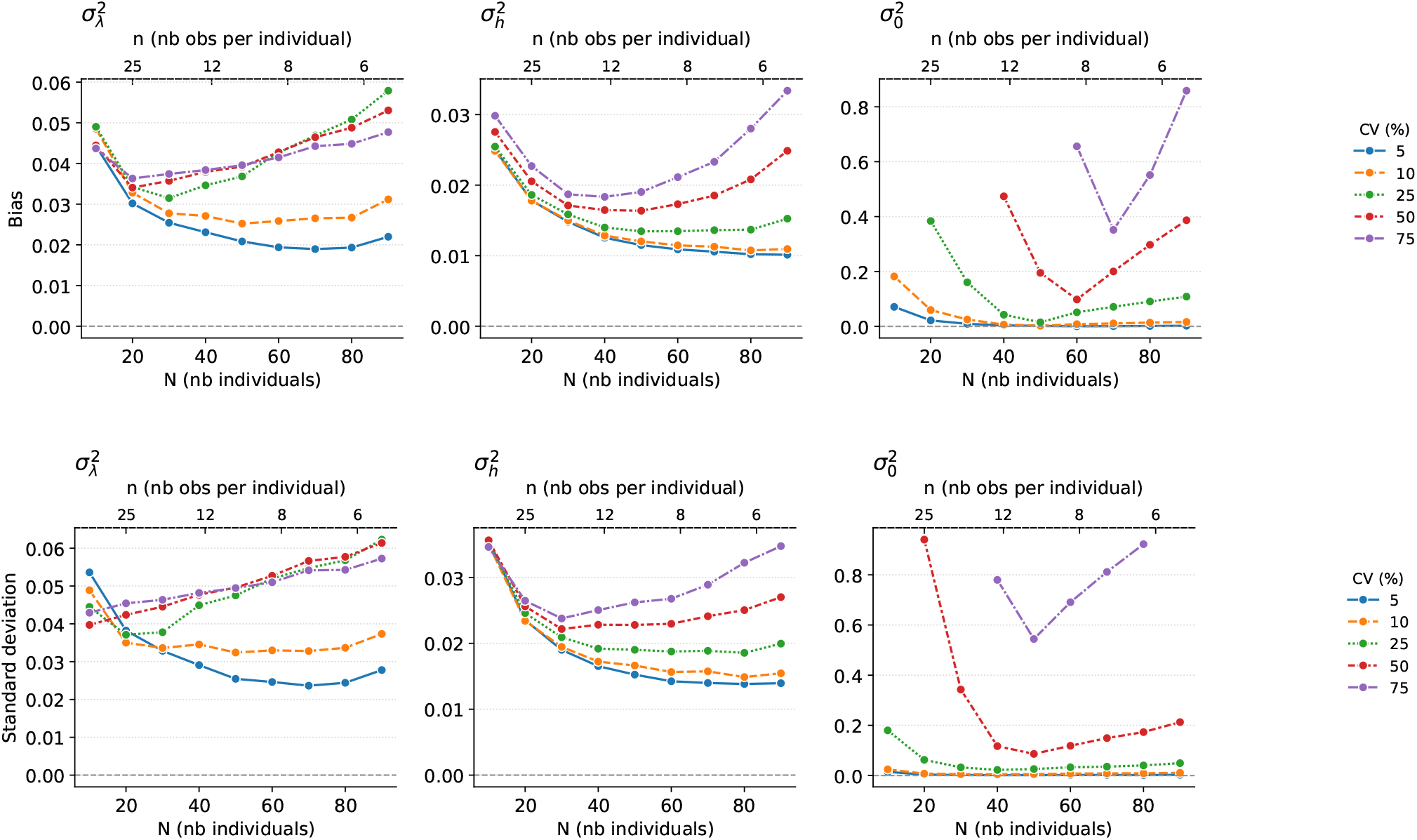
(top) Bias (in absolute value) of each variance parameter as a function of the sample structure, *i*.*e*. the number of individuals in the population (principal x-axis) and the number of observations per individual (secondary x-axis), for different coefficients of variation for the random effects and the parameter set *θ*_1_, computed over *K* = 1000 repetitions. (bottom) Standard deviation of each variance parameter as a function of the sample structure, *i*.*e*. the number of individuals in the population (principal x-axis) and the number of observations per individual (secondary x-axis), for different coefficients of variation for the residual variability and the parameter set *θ*_1_, computed over *K* = 1000 repetitions.

#### Effect of model misspecification

Fig. 11 depict results corresponding to a misspecification of the noise structure (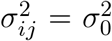 instead of 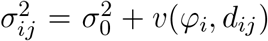). When the noise structure is misspecified, we observe an over-estimation of the residual variance term 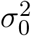 that compensates for the extra-variability of the data compared to the case where no interaction stochasticity would be present. We also observe an over-estimation of the variance component *σ*_*λ*_, in particular when the sample size *N* increases and the number of measures per individual *n* decreases. Since the estimate of residual variance is decreasing at the same time, it could be explained by the fact that the extra-variability is absorbed by *σ*_*λ*_.

**Figure 11:**
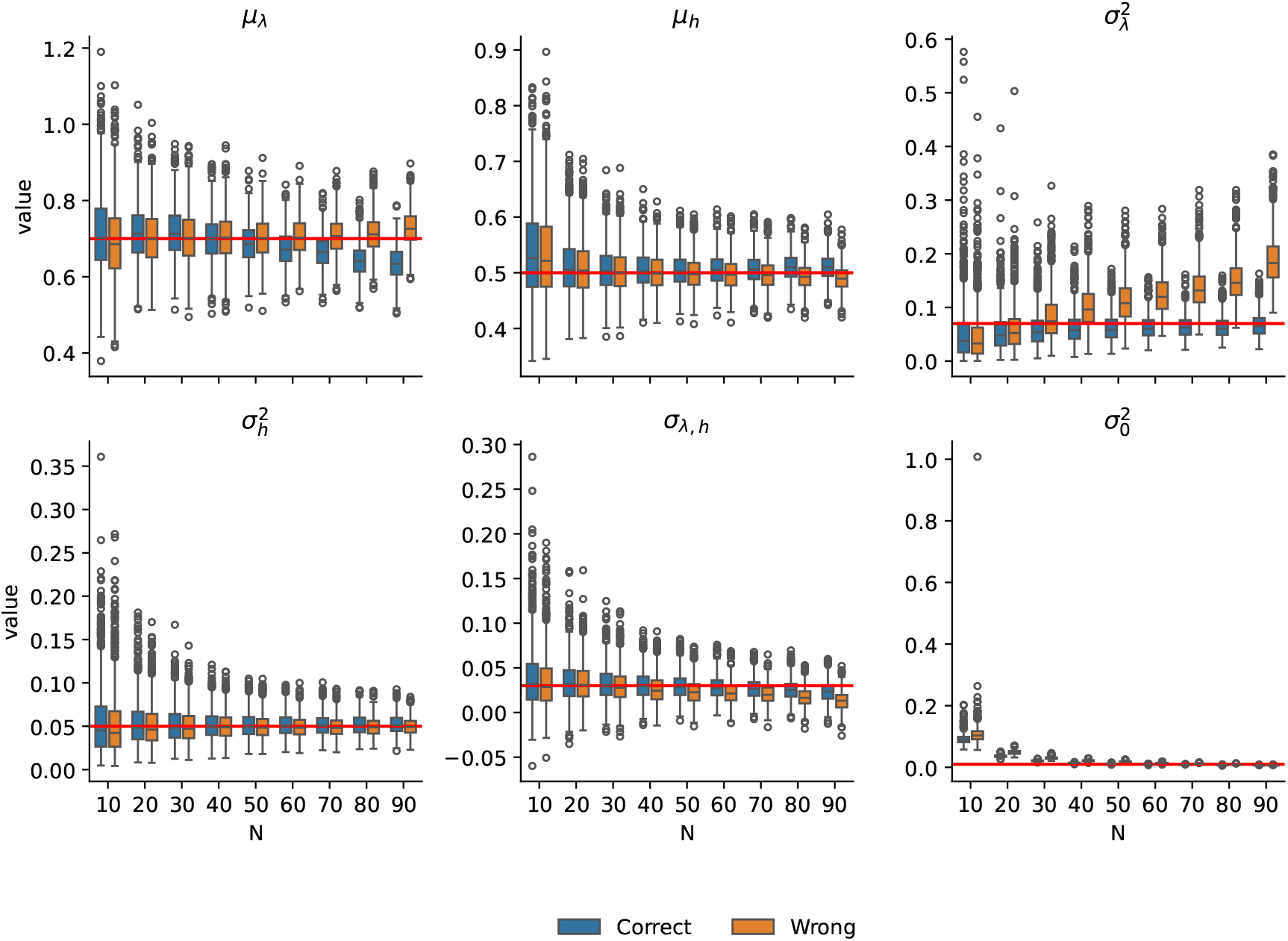
Effect of a misspecification of the noise structure on the parameter estimates, as a function of the sample size *N*. In blue, parameter estimates in the correctly specified model and in orange, parameter estimated in the misspecified model, over *K* = 1000 datasets

### 3.2 Model selection

We only focused on model selection with respect to the noise structure. Recall that the true data-generating process combines both variability from interaction stochasticity *v*(·) and residual measurement errors 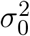. We compare this full model to a simpler alternative that includes only residual errors, without accounting for interaction stochasticity, for different signal-to-noise ratio. Simulation results are provided in Tab. 1 for different levels of random effects variability and in Tab. 2 for different levels of residual variance. They show that, regardless of the dataset structure (values of *N* and *n*), the ability to detect interaction stochasticity decreases as residual variance increases — in other words, distinguishing between mechanistic and residual sources of variation becomes increasingly difficult. We also observe that higher between-predator variability similarly reduces the ability to detect the presence of interaction stochasticity. Similar experiments were made when the random effects are assumed to be correlated, and led to similar results.

**Table 1:** Proportion of datasets for which the smallest BIC is associated to the correct model for different coefficient of variation (cv) for the random effects.

| $\begin{array}{c\vert c} & N \\ \hline cv & \end{array}$ | 10 | 20 | 30 | 40 | 50 | 60 | 70 | 80 | 90 |
| --- | --- | --- | --- | --- | --- | --- | --- | --- | --- |
| 25 | 0.56 | 0.64 | 0.83 | 0.96 | 0.99 | 1.00 | 1.00 | 1.00 | 0.99 |
| 50 | 0.47 | 0.45 | 0.61 | 0.78 | 0.83 | 0.90 | 0.92 | 0.95 | 0.92 |
| 75 | 0.44 | 0.38 | 0.39 | 0.37 | 0.36 | 0.36 | 0.34 | 0.37 | 0.32 |

**Table 2:** Proportion of datasets for which the smallest BIC is associated to the correct model for different noise to signal ratios.

| $\begin{array}{c\vert c} & N \\ \hline cv & \end{array}$ | 10 | 20 | 30 | 40 | 50 | 60 | 70 | 80 | 90 |
| --- | --- | --- | --- | --- | --- | --- | --- | --- | --- |
| 5 | 0.51 | 0.54 | 0.72 | 0.90 | 0.95 | 0.99 | 0.99 | 0.99 | 0.98 |
| 10 | 0.50 | 0.52 | 0.61 | 0.75 | 0.83 | 0.90 | 0.92 | 0.95 | 0.94 |
| 25 | 0.47 | 0.50 | 0.52 | 0.55 | 0.56 | 0.60 | 0.62 | 0.65 | 0.66 |
| 50 | 0.50 | 0.50 | 0.50 | 0.51 | 0.50 | 0.51 | 0.50 | 0.50 | 0.49 |
| 75 | 0.51 | 0.50 | 0.50 | 0.51 | 0.50 | 0.48 | 0.49 | 0.47 | 0.47 |

### 3.3 Application on a real dataset [34]

Table 3 gives the parameter estimates along with their 95% confidence intervals (CIs), with or without considering interaction stochasticity. The CI were computed using the inverse of the Fisher Information matrix, which is computed by the SGD algorithm, and the delta-method. The introduction of interaction stochasticity has an impact on the variance component estimates: *σ*_*λ*_ and *σ*_0_ are estimated a bit smaller while *σ*_*h*_ and the correlation coefficient *σ*_*λ,h*_ are estimated a bit larger in this case. Moreover, the CIs are smaller when interaction stochasticity is included in the model, especially for the variance components parameters.

**Table 3:** 95% confidence intervals for the parameters of the 3D functional model on the real data set.

| Parameter | No interaction stochasticity |  | With interaction stochasticity |  |
| --- | --- | --- | --- | --- |
|  | Estimate | 95% CI | Estimate | 95% CI |
| $\mu_\lambda$ | 0.1307 | [0.1119 ; 0.1496] | 0.1259 | [0.1058 ; 0.1460] |
| $\mu_h$ | 0.0126 | [0.0096 ; 0.0155] | 0.0124 | [0.0099 ; 0.0148] |
| $\sigma_\lambda$ | 0.0453 | [0.0280 ; 0.0626] | 0.0397 | [0.0262 ; 0.0533] |
| $\sigma_h$ | 0.0069 | [0.0058 ; 0.0079] | 0.0074 | [0.0065 ; 0.0084] |
| $\sigma_{\lambda,h}$ | 0.7751 | [0.6370 ; 0.9132] | 0.7844 | [0.7073 ; 0.8615] |
| $\sigma_0$ | 9.8380 | [9.1773 ; 10.499] | 9.7615 | [9.1238 ; 10.400] |
| BIC | 2015.10 |  | 2021.39 |  |

The residual variability *σ*_0_ is, by far, estimated to be the most important source of variation. However, care must be taken in the interpretation of the different levels of variability since the scales of the different processes are different. For example, when related to the estimated asymptotic number of prey ingested, i.e. to around 1*/µ*_*h*_ = 80, the residual variability is approximately 12% of this asymptotic response, which is comparable to the levels of residual variabilities that were tested on the simulated datasets. We computed the BIC associated with the average parameter estimates obtained over the 10 repetitions, and results are reported in Table 3. The BICs are very comparable between the two models (a difference of 6 corresponds to a difference of 3 in the log-likelihood scale). When comparing the individual BIC values obtained on the 10 repetitions for the two models, results are even more comparable, which strengthens the fact that the two models share very similar performances with respect to data fitting.

Fig. 12 shows for six randomly selected fish in the population (see complete graph in the appendix), the between-predator variability is well captured by the functional response model, which is able to reproduce the specific behavior of each individual in the population. The individual predictions obtained from the model fitted with or without considering interaction stochasticity are very similar.

**Figure 12:**
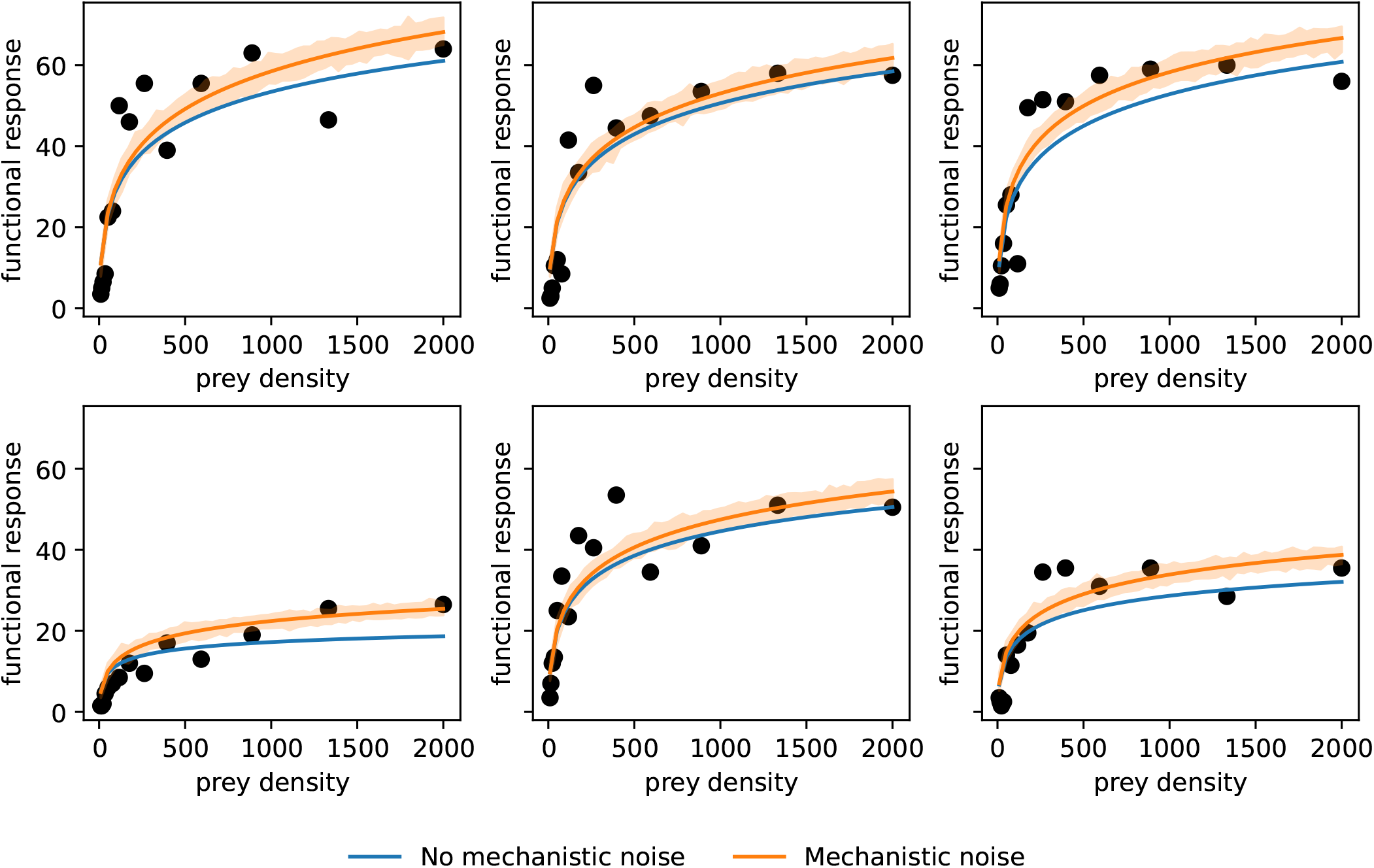
Individual predictions for six randomly selected fish in the population using the functional model run with the individual predicted values of the random effects *φ*_*i*_. The blue lines correspond to the individual predicted functional responses when no interaction stochasticity is considered, while the orange lines (resp. areas) correspond to the individual predicted functional responses when considering interaction stochasticity.

Results are very similar in both models, *i*.*e*. with or without interaction stochasticity, suggesting that between-predator variability explains a large part of the variability of the functional response. Overall, the model slightly over-estimates the upper tail of the distribution, *i*.*e*. individuals with the highest asymptotic number of prey ingested. This over-estimation is partly due to the fact that, from a prey density of approx. 800 onward, the curve of the empirical 95% quantile starts to drop, which might be explained by the small sample size. Indeed, these empirical quantiles are computed, at each observed prey density, on a total of 49 points. With a higher number of observations, we might be able to identify whether this overestimation comes from the model.

#### Comparing the One-step *vs*. two-steps estimation approaches

We estimated the model parameters log *λ*_*i*_ and log *h*_*i*_ separately for each individual curve using the `curve_fit` function from the Python `scipy` library, and then computed the empirical mean and variance of the resulting parameter estimates. Compared with the one-step mixed-effects approach (see Table 3), this procedure leads to larger estimations of all parameters: 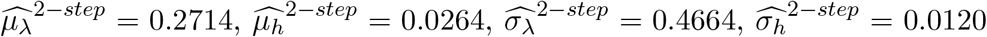. Several mechanisms can explain this discrepancy. First, the dataset contains only a small number of individuals, so the estimates are highly sensitive to those with extreme responses; their influence is not moderated by a hierarchical structure, as would be the case in mixed-effects models. Second, the curve-by-curve procedure does not account for mechanistic variance, meaning that intrinsic biological variability is mistakenly absorbed into inter-individual variability, which inflates the estimated parameters. Another important source of bias lies in the quality of the individual fits themselves. If the parameters for some curves are poorly estimated—due, for example, to noisy data, few observations, or convergence issues—these inaccurate estimates directly affect the empirical mean and variance computed across individuals. Unlike mixed-effects models, which borrow strength from the whole dataset and partially correct such errors, the separate-curve approach propagates individual misestimations to the final population-level summaries. An illustrated in a small simulation study in Section D in the appendix, the one-step approach allows for a better estimation of the inter-individual variability.

## 4 Discussion

In this study, we aimed to disentangle various sources of variation in functional response models which remains a major challenge in population ecology [38, 12, Chap. 10.2]. We developed a heteroscedastic mixed non-linear model, where both the deterministic component and the variance structure were derived from an underlying stochastic mechanistic model. This approach explicitly incorporated key sources of variation: individual differences, observational errors and model misspecification, and interaction stochasticity as a intrinsic source of variability.

### Model misspecification and estimation errors

We showed that it is possible to estimate the different sources of variability under realistic experimental conditions: with a few hundred observations, around a dozen individuals, and a limited number of parameters to estimate. We also showed that neglecting variability due to interaction stochasticity can bias estimates, particularly when the experimental design is unbalanced with respect to the number of replicates and the number of individuals (Fig. 11). This is because the model components and the sources of variation are interdependent and can exhibit compensatory behavior. Consequently, failing to account for interaction stochasticity may result in an overestimation of residual variability or a misattribution of observed variation to individual differences, ultimately distorting both parameter estimates and biological interpretations. In particular, it shows that model misspecifications regarding the variance structure, *e*.*g*. neglecting interaction stochasticity, can lead to wrong conclusions about the extent of between-predator variability. Figure 11 indeed shows that the between-individual variability in the predator searching rate, 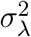, is overestimated. Figure 12 further shows that the functional response estimated for each individual can substantially differ with or without accounting for interaction stochasticity. We therefore argue that accounting for interaction stochasticity is essential when estimating the contribution of inter-individual variability to the overall variability in functional responses; otherwise, the resulting conclusions may be misleading.

As an illustration, applying our model to a real dataset resulted in the estimation of a large residual variance,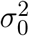. Although this variance scaled appropriately with the mean asymptotic number of prey consumed, it may also indicate that the variance from interaction stochasticity was underestimated. In theory, *σ*_0_ should capture only measurement error. However, in practice, it also absorbs model bias and misspecification, that is, the discrepancy between the proposed model and the true underlying biological process. While improving measurement procedures can help reduce observational errors, residual variability may still remain high due to an inadequate or misspecified variance structure, or from omitting intrinsic sources of variability, such as here interaction stochasticity. Evidence of such misspecification is suggested by the overestimation of upper quantiles at high prey densities in the VPC checks (Fig. 13). This pattern indicates that the number of prey consumed may plateau for predators with high consumption rates, potentially reflecting a satiety effect not accounted for in the current mechanistic model.

**Figure 13:**
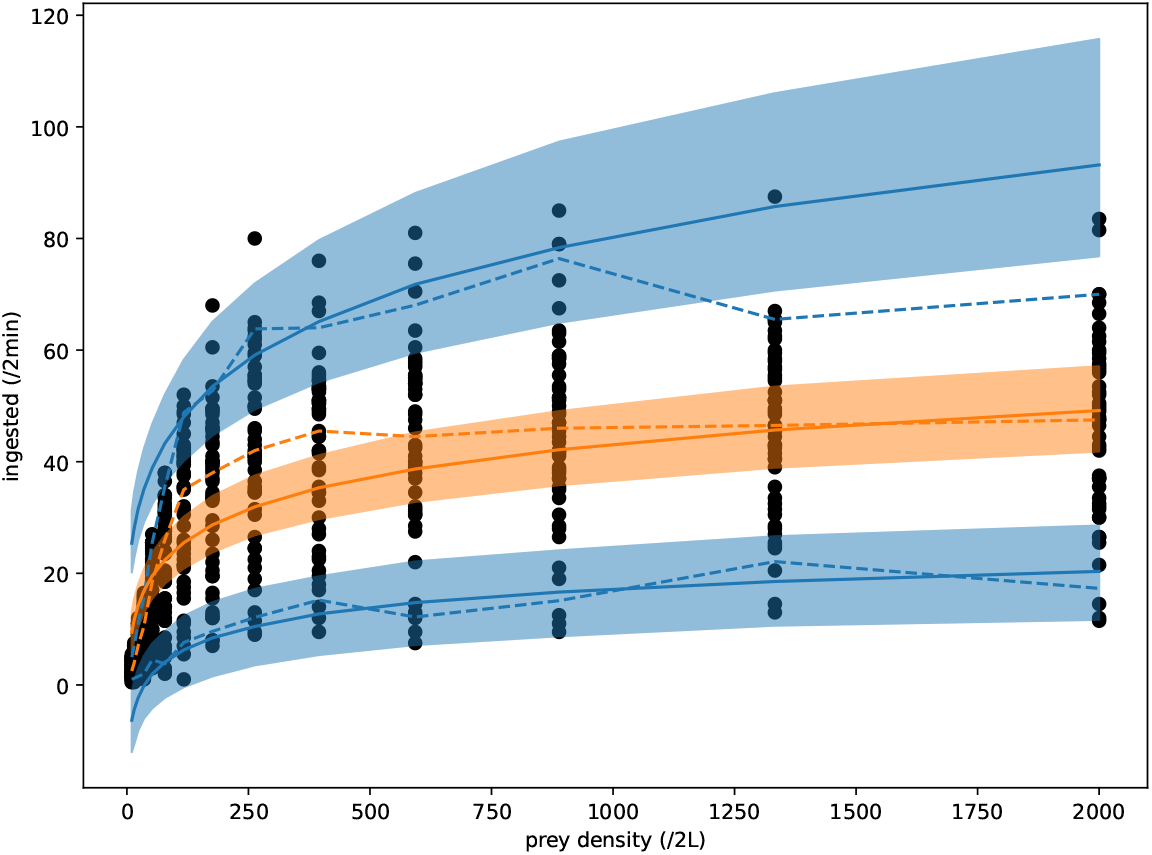
Visual Predictive Check (VPC) of the 3D functional model with interaction stochasticity on the real data set. The blue areas give the distribution of the 5% and 95% theoretical quantiles obtained from the model, while the orange are gives the distribution of the theoretical median. The solid lines are the predicted theoretical quantiles while the dotted lines are the observed quantiles. The black dots are the observed data.

### Model misspecifications regarding the variance structure

We showed that it was possible to distinguish between the residual noise and the mechanistic variability, but that it is difficult. Such a difficulty can come from potential model misspecification regarding the variance structure as the stochastic process model itself inherently limits the variability from interaction stochasticity: the mean *r* increases with *d*^1*/*3^ while the variance decreases with *d*^*−*2*/*3^. The fitting procedure may thus compensate by attributing most of the variability to measurement error, inflating *σ*_0_. This might be an issue especially in functional responses data [38, 31] as data typically show large variance. Similar estimation issues were found in [31] where strong biases were observed. In a different ecological context, [12, Chap. 10.2] conducted stochastic simulations assuming that the time between prey–predator interactions followed an exponential distribution—that is, without explicitly modeling the foraging process or accounting for spatial structure. He concluded that stochastic foraging alone could not generate the observed variability in empirical data. However, we argue that such a conclusion cannot be drawn without specifying a proper statistical model, derived from a mechanistic stochastic process, that explicitly accounts for all relevant sources of variability, as we did here. This would advocate for more appropriate stochastic models when fitting functional responses to data. For instance, [4] derived a stochastic functional response with both spatialized foraging and depletion that predicts variability to be of the same order than the mean. However, a statistical framework based on this latter model is yet to be developed, which would need much effort as the emerging distributions in [4] are unusual.

### Experimental design

We also showed that successfully disentangling the different sources of variation requires careful experimental design—specifically, balancing the trade-off between the number of individuals (*N*) and the number of observations per individual (*n*). This consideration is particularly important for accurately estimating the mean prey search speed parameter, *µ*_*λ*_, as well as the parameters associated with between-predator variability, *σ*_*λ*_ and *σ*_*h*_, especially under low signal-to-noise conditions (*i*.*e*. when *σ*_0_ is large). Given that foraging trials typically involve using each individual only once [12], our results underscore the need to reconsider conventional approaches to experimental design in this context.

### Extensions and perspectives

We focused in this study mostly on the importance of the experiment design and on comparing inference considering or not interaction stochasticity as an intrinsic mechanistic source of variation. Our statistical framework can also be used to address other issues such as i) including covariates such as individual weights or behaviors, ii) assessing the presence or importance of between-predator or between-prey variabilities in specific parameters, iii) comparing and selecting concurrent functional response models.

The standard BIC penalty may need to be adapted in those cases. In hierarchical models, the effective sample size is not always straightforward to define, especially when random effects are introduced selectively across model components. Several extensions of the BIC have been proposed to account for such structures, adjusting the penalty term to better reflect the complexity induced by individual variability. These adaptations provide more reliable guidance for model selection when testing for random effects or comparing models with different covariance structures [10, 11]. This could explain why for instance [31] found that including variability in predators’ traits did not allow improving model selection. More generally, we can hypothesize that the stochastic model used in this paper does not correctly capture the intrinsic variability. In our framework, as the distribution of the functional response is approximated from a stochastic process there is a direct mechanistic link between the mean and the variance and the parameters of the model (including the parameters describing how traits can vary between individuals). We argue that this could improve model comparison and thus help in selecting which functional response selection fits best to data.

Finally, an open question concerns the scaling parameter Δ, which arises naturally in the derivation of the stochastic process approximation and carries over into the statistical model. In our approach, we assigned Δ a fixed value determined externally by the experimental setup, making it the parameter that sets the measurement scale and provides quantitative meaning to the estimated parameters. However, one might ask whether Δ could instead be treated as an internal parameter to be estimated within the statistical framework itself. In that case, issues of identifiability would inevitably arise.

## Acknowledgements

A preprint version of this article has been peer-reviewed and recommended by PCI Ecology (https://doi.org/10.24072/pci.ecology.100808). This work was supported by the Chair ‘Modélisation Mathématique et Biodiversité’ of Veolia Environnement-Ecole Polytechnique-Museum National d’Histoire Naturelle-Fondation X, by the ANR ABIM (ANR-16-CE40-0001) and by the grant 80PRIME 2023 of the CNRS. The authors acknowledge the support of the CDP C2EMPI, as well as the French State under the France-2030 programme, the University of Lille, the Initiative of Excellence of the University of Lille, the European Metropolis of Lille for their funding and support of the R-CDP-24-004-C2EMPI project. The authors thank Vincent Bansaye, Jean-René Chazottes and Geoffroy Berthelot for helpful discussions, and Arne Schröder and Gregor Kalinkat for kindly sharing their dataset. This paper is dedicated to Elizabeta Vergu who passed away in the early stage of the current work.

## A Appendix: detailed implementations of algorithms

### A.1 Parameter estimation

To describe the algorithms, we introduce the following additional notations, which complement those used in the main body of the paper. Let *y*_*i*_ denote the vector of observations for subject *i*, where *i* = 1, …, *N*. Also, let *p*_*θ*_(·|*y*_*i*_) represent the conditional density of *φ*_*i*_ given *y*_*i*_ for a fixed value *θ* of the model parameter, and *f* (*y*_*i*_, *φ*_*i*_; *θ*) the joint density of *y*_*i*_ and *φ*_*i*_ for the parameter value *θ*. These distributions depend on model (1) specification. The two algorithms below iterate for *K* iterations, producing a sequence of estimates of the unknown parameter *θ*, denoted by (*θ*_*k*_), using a decreasing sequence of step sizes (*γ*_*k*_). Guidelines for choosing these step sizes can be found in [1] for the stochastic gradient descent and [13] for the SAEM algorithm. An important aspect to understand about the operation of these algorithms is that *φ*_*i*_ is a latent variable, meaning that we do not have an observed value for it. Consequently, it must be simulated at each iteration. It is simulated either from its exact conditional distribution *p*_*θ*_(·|*y*_*i*_) or through a small number of iterations of an MCMC procedure with kernel Π_*θ*_(·|·, *y*_*i*_), assumed to have the conditional distribution as its stationary law.

#### A.1.1 Stochastic gradient descent

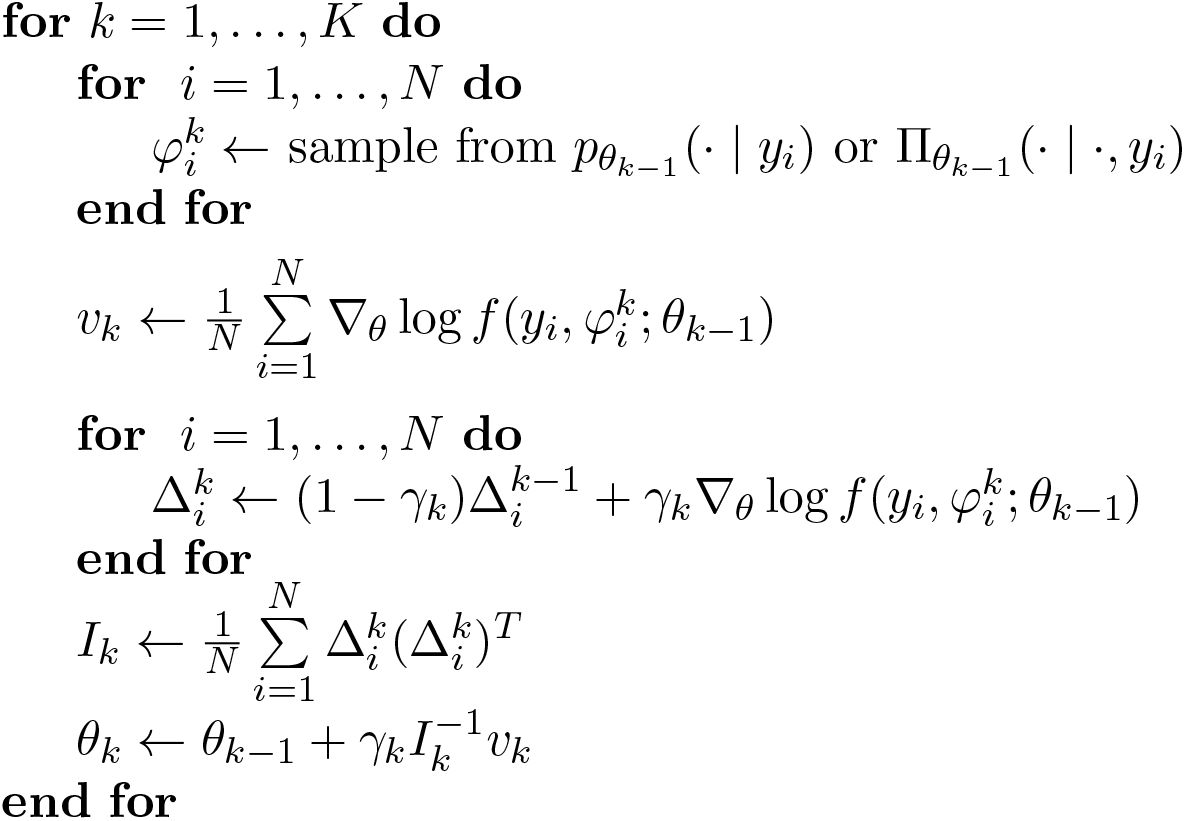

The final estimates of the Fisher information matrix and the model parameters correspond to quantities *I*_*K*_ and *θ*_*K*_ evaluated at the last iteration, assuming the algorithm has converged.

#### A.1.2 Stochastic approximation EM (SAEM) algorithm

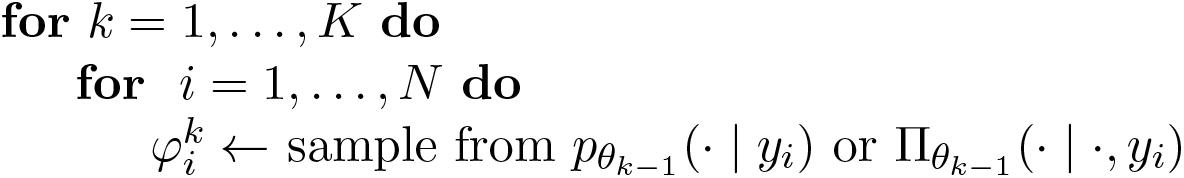
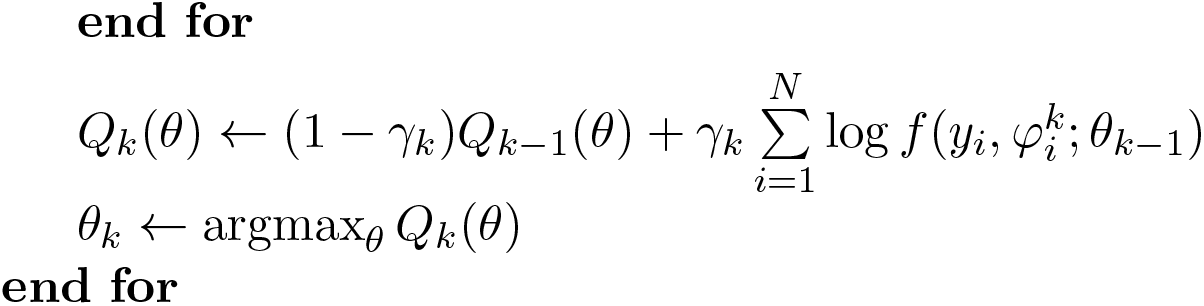

Note that for this algorithm, the implementation of an iteration is significantly simplified when the model belongs to the curved exponential family (see [20]).

The final estimate of the model parameters correspond to *θ*_*K*_ evaluated at the last iteration, assuming the algorithm has converged.

### A.2 Log-likelihood estimation

To estimate the log-likelihood in a nonlinear mixed-effects model, we need to evaluate integrals over the random effects. The marginal likelihood is given by:

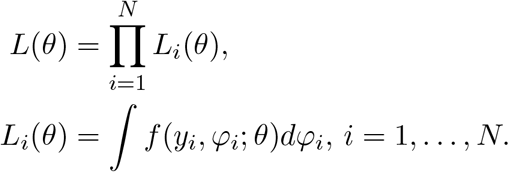

These integrals are generally intractable. We can use Monte-Carlo methods such as Importance Sampling to approximate them. The idea is to draw many samples of the random effects, 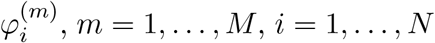, *m* = 1, …, *M, i* = 1, …, *N*, from a known, easy-to-sample distribution 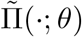 and then weight each sample according to the following formula:

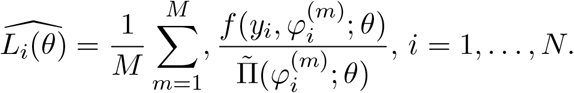

The log-likelihood is then approximated by

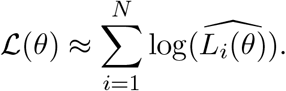

A critical aspect of this method is the choice of the importance distribution 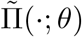, which should be close enough to the conditional distribution of the random effects given the observations to ensure estimation stability. In practice, there is no universal choice: selecting a good importance distribution often involves trial and error, along with stability checks.

### B Convergence of the SGD algorithm on the real dataset

**Figure A.1:**
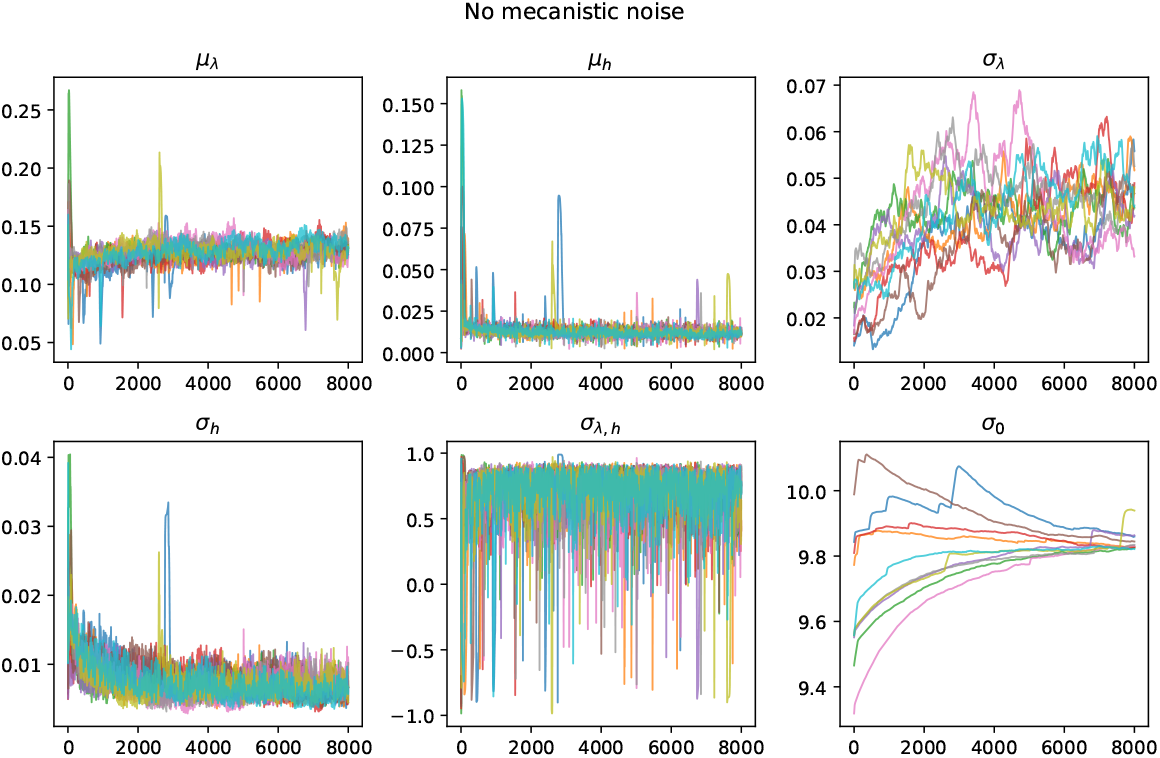
Evolution of the parameter estimates along the iterations of the SGD algorithms, for each of the ten runs corresponding to ten different initializations, when no mechanistic noise is included in the model

**Figure A.2:**
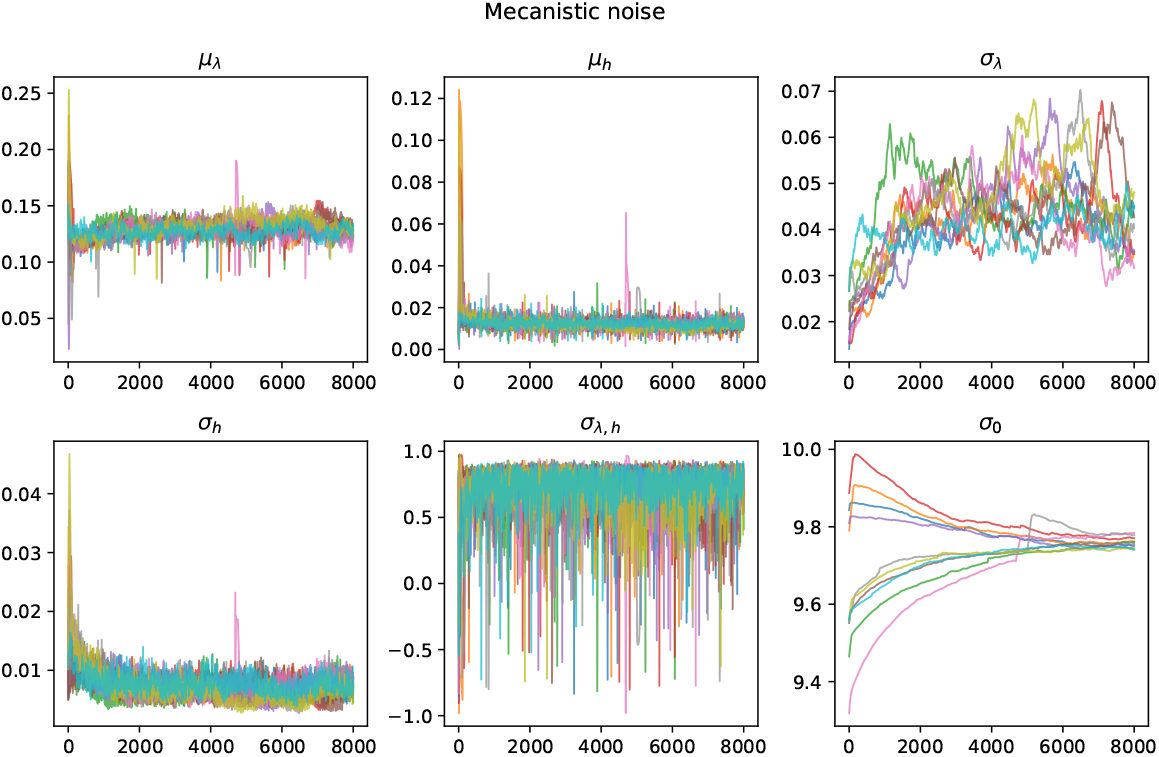
Evolution of the parameter estimates along the iterations of the SGD algorithms, for each of the ten runs corresponding to ten different initializations, when mechanistic noise is included in the model

### C Individual predictions on the real dataset

**Figure A.3:**
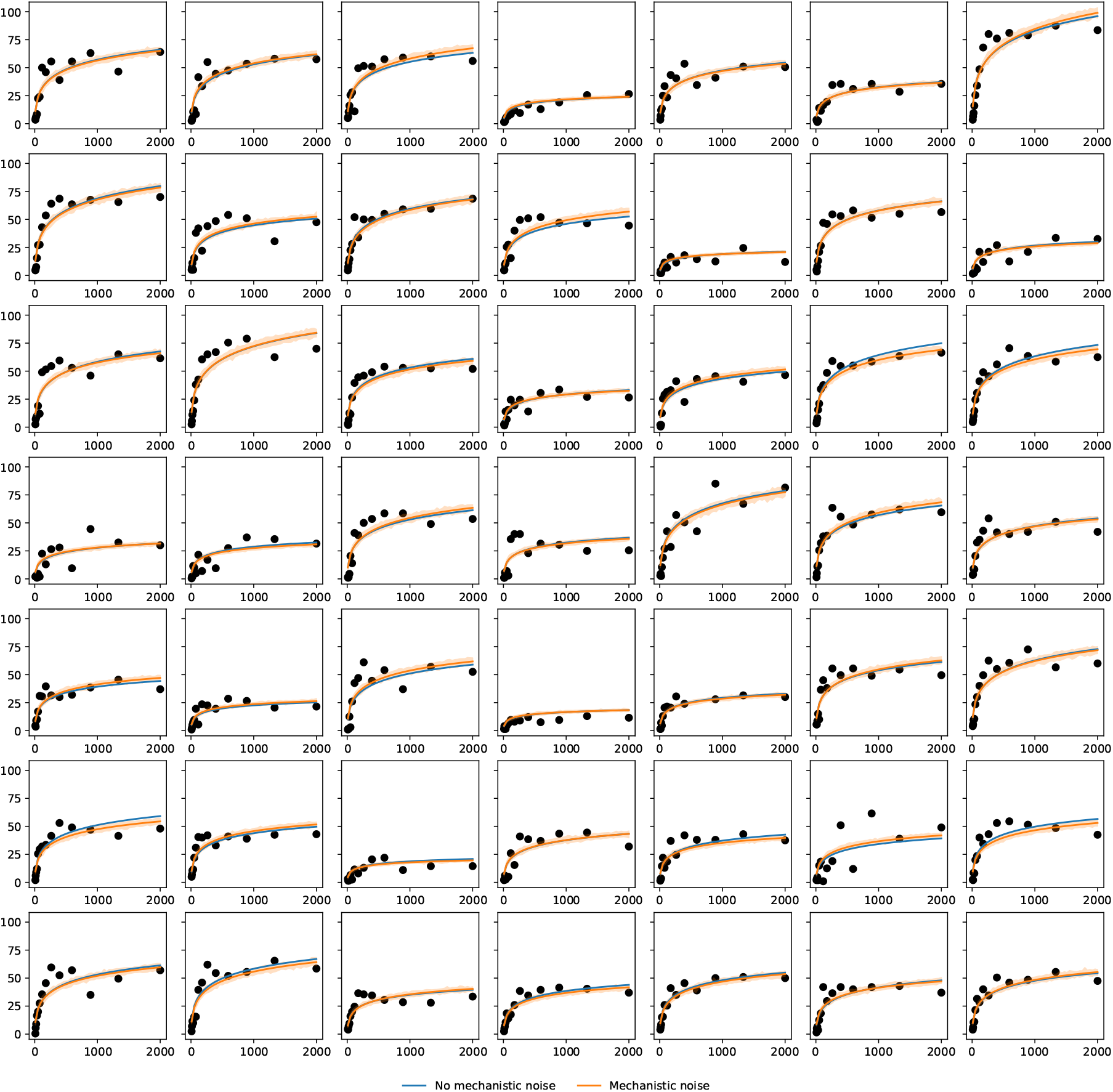
Individual predictions for all the population using the functional model run with the individual predicted values of the random effects *φ*_*i*_.

### D Advantages of the mixed-model approach compared to a two-stage estimation

We illustrate the difference between the one-step and the two-step approaches on a simulation study. In the former, the parameters associated to the different sources of variability are estimated jointly in a single step. This is the mixed-model approach that we adopted in this paper. On the contrary, the latter first proceeds with a separate estimation of the functional response parameters for each individual of the population, before assessing their between-individual variability in a second step.

We simulated the functional responses of *N* individuals using the model in Eq (1), but without mechanistic noise (*i*.*e*. 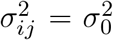), at *n* different prey densities. For the two-step approach, we fitted each individual curve separately, to obtain individual estimates for *λ* and *h* denoted by 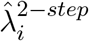 and 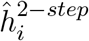. We then estimated the mean and standard-deviations of these individual parameters, denoted by 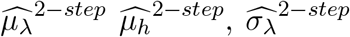 and 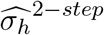 using the empirical means and standard-deviations of the individual estimates. For the one-step approach, we fitted a nonlinear mixed-effects model to all the population, with independent random effects for *λ* and *h* and used the estimated means and standard deviation of these random effects for 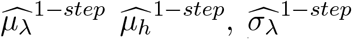 and 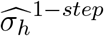. We also used the individual predictions of the random effects as 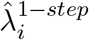 and 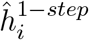.

We compared the population estimates to the real values used to generate the data, for several combinations of (*N, n*) ∈ {(10, 10), (10, 50), (50, 10), (100, 5), (100, 10)} by simulated a total of *M* = 1000 datasets. Results are given in Fig. A.4

Overall, we can see that the one-step approach performs better globally in every setting. When the sample size is small (*N* = 10), the one-step approach exhibits a higher variance, due to the fact that the theoretical guarantees of this method requires that the number of individuals *N* is large enough. When the number of measurements per individuals is small, the two-step approach exhibits a high bias in the standard-deviation estimates. This is particularly true when *n* = 5 (the blue boxplot). This trend is not counter-balanced by an increased number of individuals (see for example the persistent bias when *n* = 10, in boxplots 1, 3 and 5 in Fig. A.4).

**Figure A.4:**
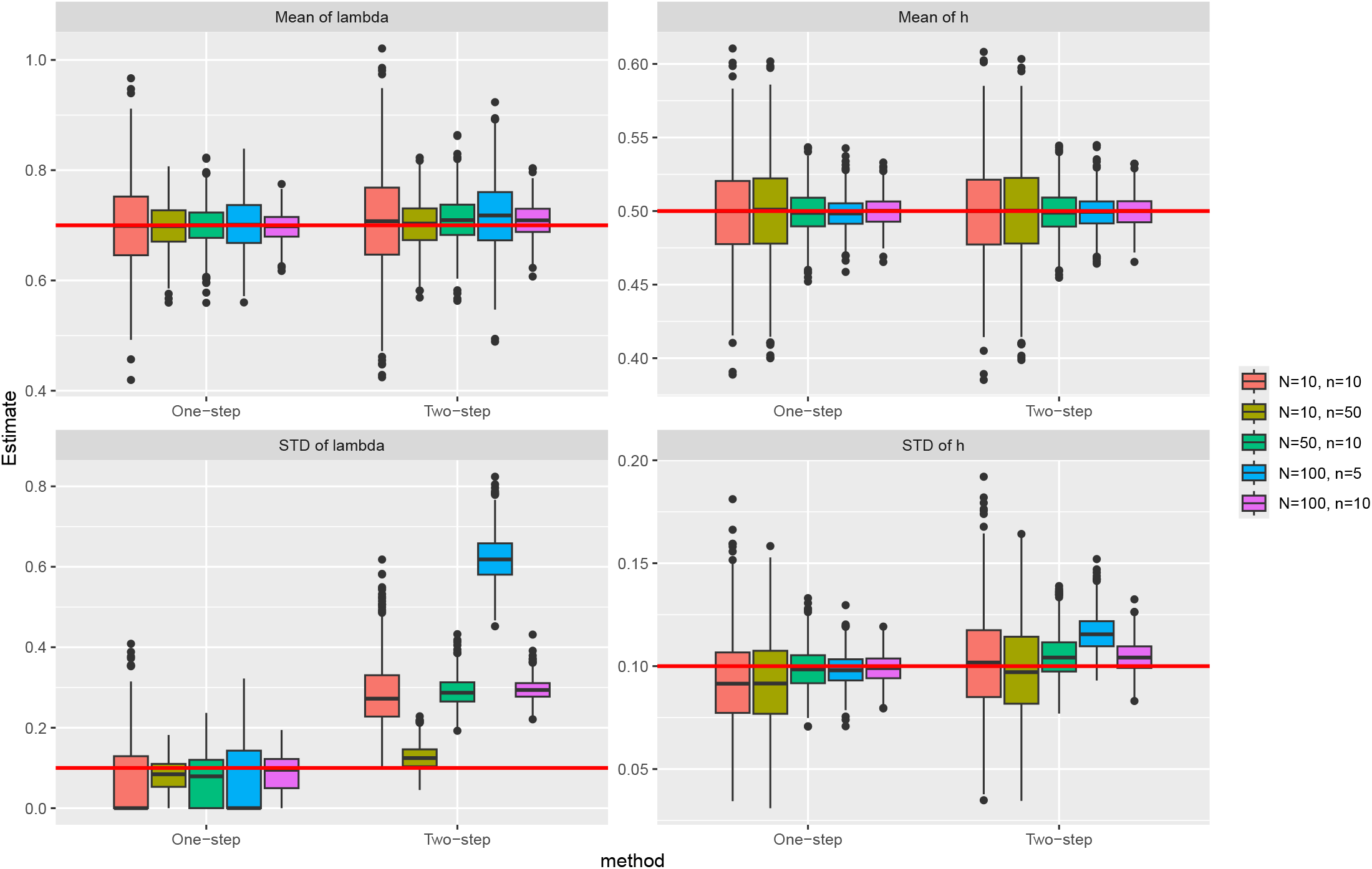
Comparison of the estimates of the mean and standard deviation of the individual parameters obtained with a one-step or a two-step procedure.

